# Macrophage-to-sensory neuron crosstalk mediated by Angiotensin II type-2 receptor elicits neuropathic pain

**DOI:** 10.1101/166546

**Authors:** Andrew J. Shepherd, Aaron D. Mickle, Bryan A. Copits, Páll Karlsson, Suraj Kadunganattil, Judith P. Golden, Satya M. Tadinada, Madison R. Mack, Simon Haroutounian, Annette D. de Kloet, Vijay K. Samineni, Manouela V. Valtcheva, Lisa A. McIlvried, Tayler D. Sheahan, Sanjay Jain, Pradipta R. Ray, Yuriy M. Usachev, Gregory Dussor, Brian S. Kim, Eric G. Krause, Theodore J. Price, Robert W. Gereau, Durga P. Mohapatra

**Affiliations:** Department of Anesthesiology and Washington University Pain Center, Washington University School of Medicine in St. Louis, MO, 63110, USA; Department of Pharmacology, The University of Iowa Carver College of Medicine, Iowa City, IA, 52242, USA; Danish Pain Research Center, Department of Clinical Medicine, Aarhus University Hospital, DK-8000 Aarhus C, Denmark; Department of Clinical Medicine – Core Center for Molecular Morphology, Section for Stereology and Microscopy, Aarhus University Hospital, DK-8000 Aarhus C, Denmark; Department of Dermatology and Center for the Study of Itch, Washington University School of Medicine in St. Louis, MO, 63110, USA; Department of Physiology and Functional Genomics, College of Medicine, University of Florida, Gainesville, FL, 32610, USA; Departments of Medicine, Pathology and Immunology, Washington University School of Medicine in St. Louis, MO, 63110, USA; School of Behavioral and Brain Sciences, University of Texas at Dallas, Richardson, TX, 75080, USA; Department of Pharmacodynamics, College of Pharmacy, University of Florida, Gainesville, FL, 32610, USA; Department of Neuroscience, Washington University School of Medicine in St. Louis, MO, 63110, USA; Center for the Investigation of Membrane Excitability Diseases, Washington University School of Medicine in St. Louis, MO, 63110, USA; Siteman Cancer Center, Washington University School of Medicine in St. Louis, MO, 63110, USA

## Abstract

Peripheral nerve damage initiates a complex series of cellular and structural processes that culminate in chronic neuropathic pain. Our study defines local angiotensin signaling via activation of the Angiotensin II (Ang II) type-2 receptor (AT2R) on macrophages as the critical trigger of neuropathic pain. An AT2R-selective antagonist attenuates neuropathic, but not inflammatory pain hypersensitivity in mice, and requires the cell damage-sensing ion channel transient receptor potential family-A member-1 (TRPA1). Mechanical and cold pain hypersensitivity that are characteristic of neuropathic conditions can be attenuated by chemogenetic depletion of peripheral macrophages and AT2R-null hematopoietic cell transplantation. Our findings show no AT2R expression in mouse or human sensory neurons, rather AT2R expression and activation in macrophages triggers production of reactive oxygen/nitrogen species, which trans-activate TRPA1 on sensory neurons. Our study defines the precise neuro-immune crosstalk underlying nociceptor sensitization at the site of nerve injury. This form of cell-to-cell signaling represents a critical peripheral mechanism for chronic neuropathic pain, and therefore identifies multiple analgesic targets.

## INTRODUCTION

Neuropathic pain is caused by a disease or lesion affecting the sensory nerves. Often intractable in nature, neuropathic pain serves no protective biological function, and is estimated to affect ∼3-17% of the global population (1). The etiology of neuropathic pain is complex, therefore presenting a formidable challenge to its effective management. It is closely associated with the use of cancer chemotherapeutic drugs, diabetic neuropathy, post-herpetic neuralgia (PHN), traumatic injury and trigeminal neuralgia. The lack of a precise mechanistic understanding has undoubtedly hampered the development of effective analgesics for neuropathic pain, which is poorly managed by existing drugs (2, 3). However, Angiotensin II (Ang II) type-2 receptor (AT2R) antagonists have recently proven efficacious in preclinical models of neuropathic and cancer pain (4), and the AT2R antagonist EMA401 has shown effective pain relief in PHN patients in phase II clinical trials (5). Importantly, the site of action for this antagonist remains controversial.

The major effector of the renin-angiotensin system (RAS), Ang II is generated from Angiotensinogen (Agt) and Angiotensin I by renin and angiotensin converting enzyme (ACE), respectively. Ang II regulation of blood pressure has been well-established, with most of its physiological actions ascribed to type 1 (AT1R) receptor signaling; however, the role of AT2R has remained enigmatic (6). Expression of AT2R in the brain has been reported, contributing to a number of functions such as regulation of drinking behavior and motor activity (6-8). Also, recent findings have suggested that AT2R in peripheral sensory neurons is involved in pain modulation (4). Specifically, signaling originating from Gα_s_-coupled AT2R in sensory neurons was shown to elicit peripheral pain sensitization (9, 10). In contrast, a myolactone toxin from Buruli ulcer-causing bacteria was shown to activate Gα_i/o_-coupled AT2R in sensory neurons, leading to analgesia in mice (11). Interestingly, a recent follow-up study demonstrated that the AT2R antagonist EMA401 was unable to prevent the myolactone toxin’s effect on sensory neurons *in vitro* (12). This raises the possibility that the analgesic actions of EMA401 could result from targeting non-neuronal AT2R, or could be entirely independent of AT2R antagonism. More recently, it was proposed that the AT2R competitive antagonist PD123319 (and EMA401) could indirectly increase levels of the Ang II-cleavage product Ang1-7, which activates the Mas1 receptor to elicit anti-nociceptive effects in a rodent model of bone cancer pain (13). Therefore, establishing the mechanistic underpinnings of angiotensin signaling in diverse chronic pain states is essential for further therapeutic developments.

Using the spared nerve injury (SNI) model of neuropathy and the complete Freund’s adjuvant (CFA) model of inflammation in mice, we corroborate the effectiveness of AT2R antagonism selectively in neuropathic pain hypersensitivity. Elevated levels of Ang II are detected specifically in injured sciatic nerves, and Ang II acutely induces tactile hypersensitivity, similar to that associated with neuropathy. By combining pharmacological and genetic manipulation, we show the requirement of AT2R and the cold/mechano-sensitive pain receptor TRPA1 in Ang II- and neuropathy-induced pain hypersensitivity. However, our in-depth investigation shows no AT2R expression in mouse or human sensory neurons, and Ang II does not directly influence sensory neuron function. Instead, our study describes the critical role of macrophages (MΦs) and AT2R expression therein at the site of nerve injury in the development of Ang II- and neuropathy-induced pain hypersensitivity. Furthermore, we identify Ang II-AT2R-mediated reactive oxygen/nitrogen species (ROS/RNS) production in MΦs as the critical trigger of TRPA1 activation on sensory neurons. Our findings comprehensively define the role of angiotensin signaling and intercellular redox communication between MΦs and sensory nerves in the development of chronic pain resulting from neuropathy.

## RESULTS

### Angiotensin II – AT2R activation is critical for nerve injury-induced mechanical pain hypersensitivity

We began with an unambiguous verification of the critical role of AT1R and AT2R antagonism in alleviating nerve injury-induced chronic pain. SNI-induced peripheral neuropathy in mice elicited long-lasting mechanical hypersensitivity (Figure 1 A and B). Instead of simply determining mechanical paw withdrawal threshold, the magnitude of total mechanical sensitivity on mouse hindpaws in response to increasing von Frey filament strength was determined, as detailed earlier in several reports, and outlined in Supplemental Figure 1A. Systemic administration of the AT2R antagonist PD123319 dose-dependently attenuated SNI-induced mechanical hypersensitivity in male and female mice to a similar extent (Figure 1A, and Supplemental Figure 1 B and D). However, administration of the AT1R antagonist losartan did not influence SNI-induced mechanical hypersensitivity (Figure 1C). Administration of PD123319 or losartan alone in sham mice did not alter hindpaw mechanical sensitivity, and no change in mechanical sensitivity was observed in the contralateral hindpaws of SNI mice (Figure 1 B and C, and Supplemental Figure 1C). SNI did not influence hindpaw heat sensitivity in male or female mice (Supplemental Figure 1 E and F), as demonstrated previously (14, 15). We next verified whether AT2R inhibition of SNI-induced mechanical hypersensitivity operates at a central or peripheral level. Intrathecal (i.t.) administration of PD123319 did not attenuate mechanical hypersensitivity (Figure 1D), but peri-sciatic delivery, as with systemic administration (i.p.), proved effective (Figure 1E). Attenuation of SNI-induced mechanical hypersensitivity by PD123319 was independent of any hemodynamic changes, since PD123319 administration, unlike losartan, did not influence blood pressure in mice (Supplemental Figure 2A). Vascular permeability of the hindpaw was also unaffected by PD123319, as determined by Evans Blue extravasation (Supplemental Figure 2B). Furthermore, we observed that systemic administration of PD123319 did not attenuate mechanical and heat hypersensitivity due to chronic hindpaw inflammation induced by CFA (Figure 1F and Supplemental Figure 3).

**Figure 1:**
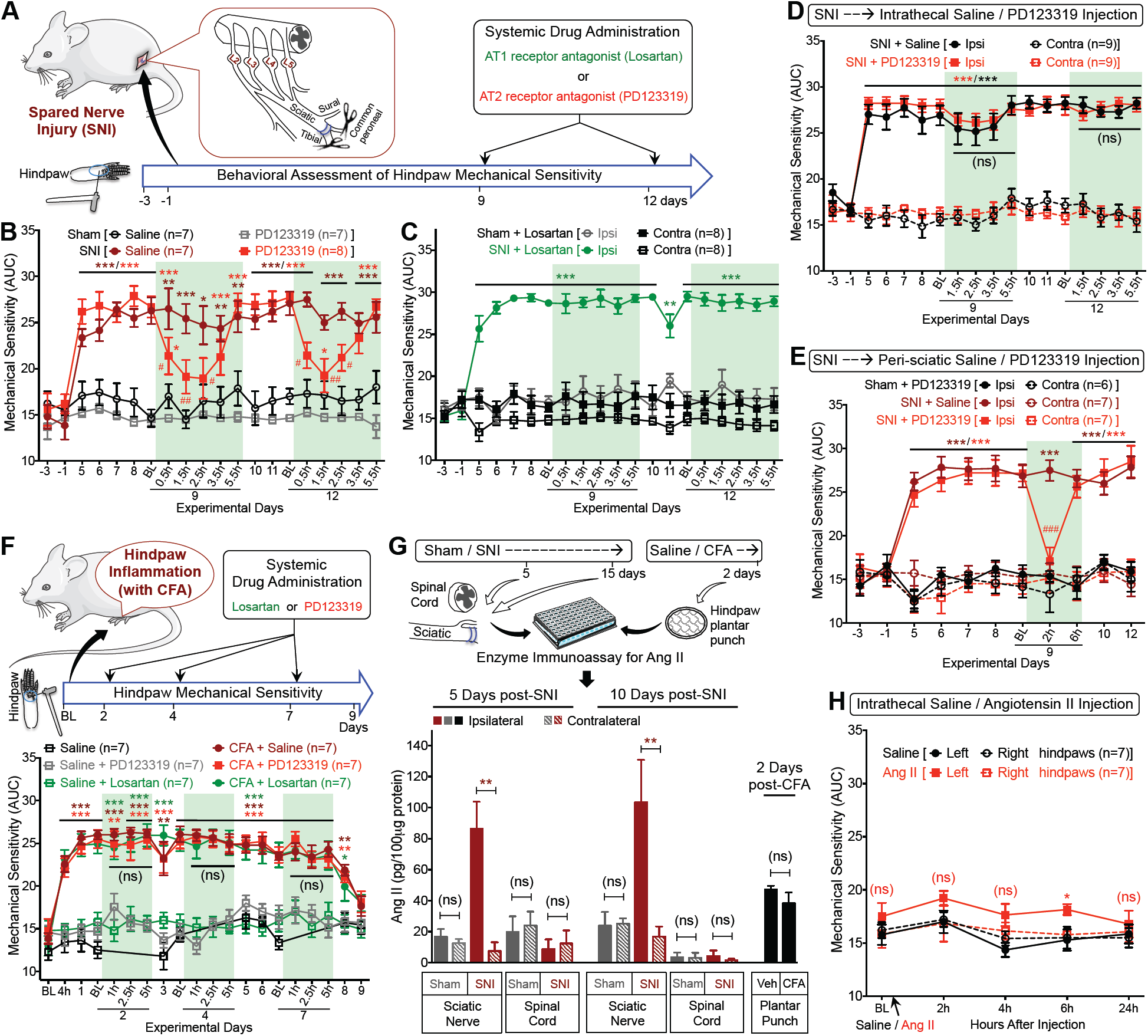
Peripheral AT2R activation mediates neuropathic pain hypersensitivity. (**A**) Experimental scheme depicting nerve injury-induced neuropathy, pain behavioral assessments, and drug administration timeline in C57BL/6 (B6) mice for **B-E** (data analysis scheme in fig. S1A). (**B**) Systemic administration of PD123319 (10 mg/kg; i.p.) attenuates SNI-induced mechanical hypersensitivity. Mean ± SEM; \**p*<0.05, \*\**p*<0.01 and ^***^*p*<0.001, versus sham+saline/PD123319 groups; ^#^*p*<0.05 and ^##^*p*<0.01, versus SNI+saline group. (**C**) Losartan (10 mg/kg; i.p.) has no effects on SNI-induced mechanical hypersensitivity. Mean ± EM; \*\**p*<0.01 and ^***^*p*<0.001, versus sham+losartan-ipsi group. (**D**) Intrathecal PD123319 (10 mg/kg) does not attenuate SNI-induced mechanical hypersensitivity. Mean ± SEM; ^***^*p*<0.001, versus contra groups; not significant (ns), versus SNI+saline-ipsi group. (**E**) Peri-sciatic PD123319 administration (10 mg/kg) attenuates SNI-induced mechanical hypersensitivity. Mean ± SEM; ^***^*p*<0.001, versus contra groups; ^###^*p*<0.001, versus SNI+saline-ipsi group. (**F**) PD123319 or losartan (10 mg/kg for each; i.p.) does not attenuate peripheral inflammation (CFA)-induced mechanical hypersensitivity, as detailed in the experimental scheme shown at the top. Mean ± SEM; \**p*<0.05, \*\**p*<0.01 and ^***^*p*<0.001, versus saline+PD123319 or saline+losartan groups. (**G**) SNI elevates Ang II levels in injured mouse sciatic nerve, but not in the spinal cord. CFA did not increase plantar hindpaw tissue Ang II levels. Mean ± SEM (n= duplicate tissue samples from 3 mice/group). \*\**p*<0.01, versus respective SNI-contralateral groups; not significant (ns), versus sham/SNI-contralateral or vehicle groups. (**H**) Intrathecal Ang II (100 pmol) does not induce significant long-lasting mechanical hypersensitivity. Mean ± SEM; \**p*<0.05 and not significant (ns), versus saline-ipsi group. Rectangular boxes in panels **B-F** denote post-drug administration behavioral assessment time points.

We next investigated if SNI was associated with changes in Ang II production. Ang II levels were elevated in the ipsilateral sciatic nerve from SNI mice, but not in contralateral or sham-operated mice. No elevation in Ang II levels was observed in the spinal cord of SNI-versus sham-operated mice, or in the plantar hindpaw skin of CFA versus saline-injected mice (Figure 1G). We also observed that Ang II (i.t.) injection did not induce mechanical hypersensitivity (Figure 1H), as reported previously (16); collectively suggesting that SNI elevates levels of Ang II in the sciatic afferent fibers, which then presumably acts on peripheral AT2R to induce mechanical hypersensitivity. Accordingly, Ang II injection into mouse hindpaws dose-dependently induced mechanical hypersensitivity, without inducing any heat hypersensitivity (Figure 2A, and Supplemental Figure 4 A and B). These effects were similar in both male and female mice, suggesting no sex differences in Ang II-induced mechanical hypersensitivity (Supplemental Figure 4 C and D). Furthermore, Ang II-induced mechanical hypersensitivity was not influenced by losartan co-administration in WT mice or Ang II injection in *Agtr1-KO* mice (Figure 1J, and Supplemental Figure 4 E and G). However, Ang II-induced mechanical hypersensitivity was completely attenuated by PD123319 co-administration in WT mice, and was absent upon Ang II injection in *Agtr2-KO* mice (Figure 1K and Supplemental Figure 4 F and G). Bradykinin injection into *Agtr2-KO* mouse hindpaws induced mechanical hypersensitivity (Supplemental Figure 4H), suggesting an absence of any gross deficits in inflammatory mediator-induced pain hypersensitivity in mice lacking functional AT2R.

**Figure 2:**
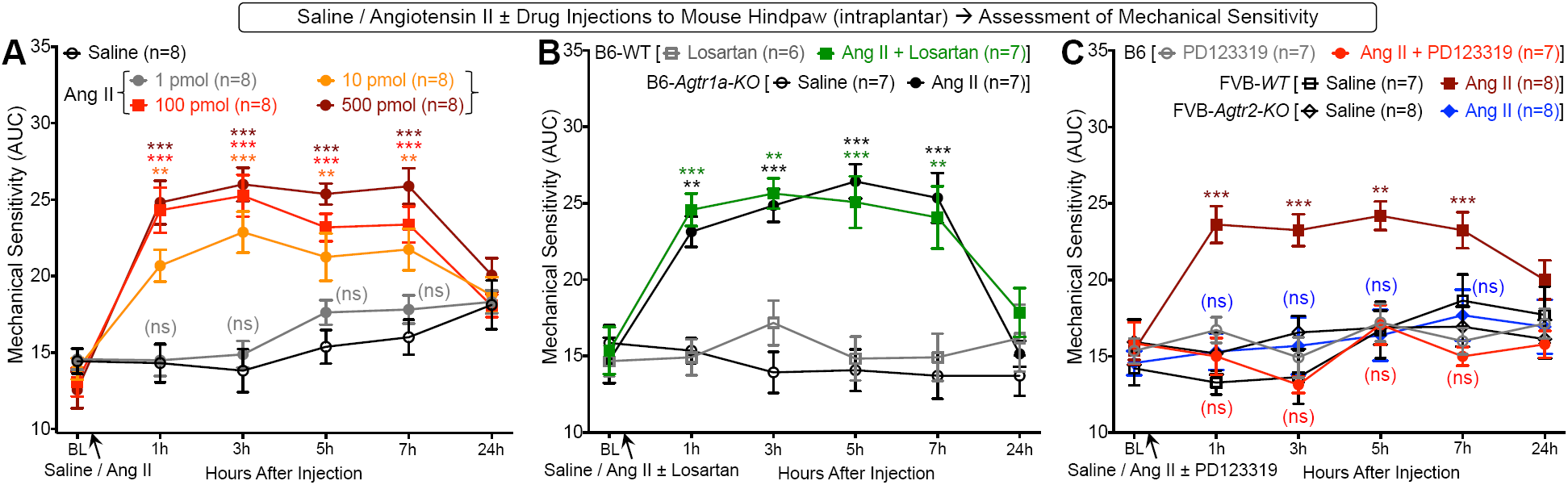
Ang II elicits mechanical hypersensitivity in mice, and is dependent on AT2R. (**A**) Ang II injection (i.pl.) dose-dependently induces hindpaw mechanical hypersensitivity in B6 mice. Mean ± SEM; ^**^*p*<0.01, ^***^*p*<0.001 and not significant (ns), versus saline group. (**B**) Ang II-induced (100 pmol, i.pl.) mechanical hypersensitivity is not attenuated by losartan co-administration (10 pmol; i.pl.) in B6-*WT* mice. Ang II (100 pmol, i.pl.) induces mechanical hypersensitivity in B6-*WT* and B6-*Agtr1a-KO* mice to a similar magnitude. Mean ± SEM; ^**^*p*<0.01 and ^***^*p*<0.001, versus B6-*WT*-losartan and B6-*Agtr1a-KO*-saline groups. (**C**) PD123319 co-administration (10 pmol; i.pl.) completely attenuates Ang II-induced (100 pmol, i.pl.) mechanical hypersensitivity in B6 mice. Ang II (100 pmol, i.pl.) induces mechanical hypersensitivity in B6-*WT* and FVB-*Agtr2-WT* mice to a similar magnitude, which is absent in FVB-*Agtr2-KO* mice. Mean ± SEM; ^*^*p*<0.05, ^**^*p*<0.01 and ^***^*p*<0.001 versus saline injection in FVB-*Agtr2-WT* mice; not significant (ns), versus FVB-*Agtr2-KO*-saline and B6-*WT*-PD123319 groups.

### AT2R and TRPA1 are essential for Ang II/neuropathy-related mechanical and cold pain hypersensitivity

To further determine the sensory receptor(s) involved in SNI/Ang II-induced mechanical hypersensitivity, we directed our attention to critical TRP channels involved in pain hypersensitivity with injury and inflammatory conditions, such as TRPA1, TRPV1 and TRPV4. SNI- and Ang II-induced mechanical hypersensitivity were completely attenuated by systemic administration of TRPA1 inhibitors A967079 and AP-18, respectively (Figure 3 A and B, and Supplemental Figure 5 A and B). Also, hindpaw injection of Ang II failed to induce mechanical hypersensitivity in *Trpa1-KO* mice (Figure 3B). In contrast, the TRPV1 inhibitor AMG9810 did not attenuate SNI-induced mechanical hypersensitivity (Figure 3A, and Supplemental Figure 5A). Furthermore, hindpaw injection of Ang II elicited mechanical hypersensitivity in both *Trpv1-KO* and *Trpv4-KO* mice to an extent similar to that observed in *WT* mice (Figure 3C), without any influence on heat hypersensitivity (Supplemental Figure 5C). Given that neuropathic conditions elicit pronounced cold hypersensitivity, and TRPA1 is known to participate in the development of cold hypersensitivity (17, 18), we assessed hindpaw cold sensitivity in SNI- and sham-operated mice (Figure 3D). SNI induced significant cold hypersensitivity, which was transiently reversed by systemic administration (i.p.) of AT2R antagonist PD123319 (Figure 3E) or TRPA1 inhibitor A967079 (Figure 3F). In order to assess a voluntary, non-reflexive measure of mechanical hypersensitivity, we subjected mice to a mechanical conflict-avoidance (MCA) assay (Figure 4A) (19). At a spike height of 5 mm in the conflict chamber, SNI mice showed a significant increase in the latency to escape from the lit chamber, which was completely attenuated upon systemic administration (i.p.) of PD123319 (Figure 4B) or A967079 (Figure 4C). Similarly, in separate cohorts of mice without prior exposure/baseline in MCA, systemic administration (i.p.) of PD123319 led to attenuation of the SNI-induced increase in the latency to escape from the lit chamber (Supplemental Figure 5D). We also performed a thermal place preference/avoidance assay to better interrogate voluntary cold hypersensitivity behavior associated with SNI (Figure 4D). Eight days post-surgery, SNI mice exhibit significant aversion to the 20^°^C plate relative to the 30^°^C plate, an effect that persisted at post-operative day 11, and was significantly attenuated by systemic administration (i.p.) of PD123319 (Figure 4E), and was subsequently reversed 24 hours later, at 12 days post-surgery (Supplemental Figure 5E). Similar inhibition of SNI-induced cold aversion was also observed at post-operative day 14 with A967079 (30 mg/Kg, i.p.; Figure 4F), and at day 16 with systemic administration (i.p.) of buprenorphine, as a pain-related behavioral validation control (Supplemental Figure 5F).

**Figure 3:**
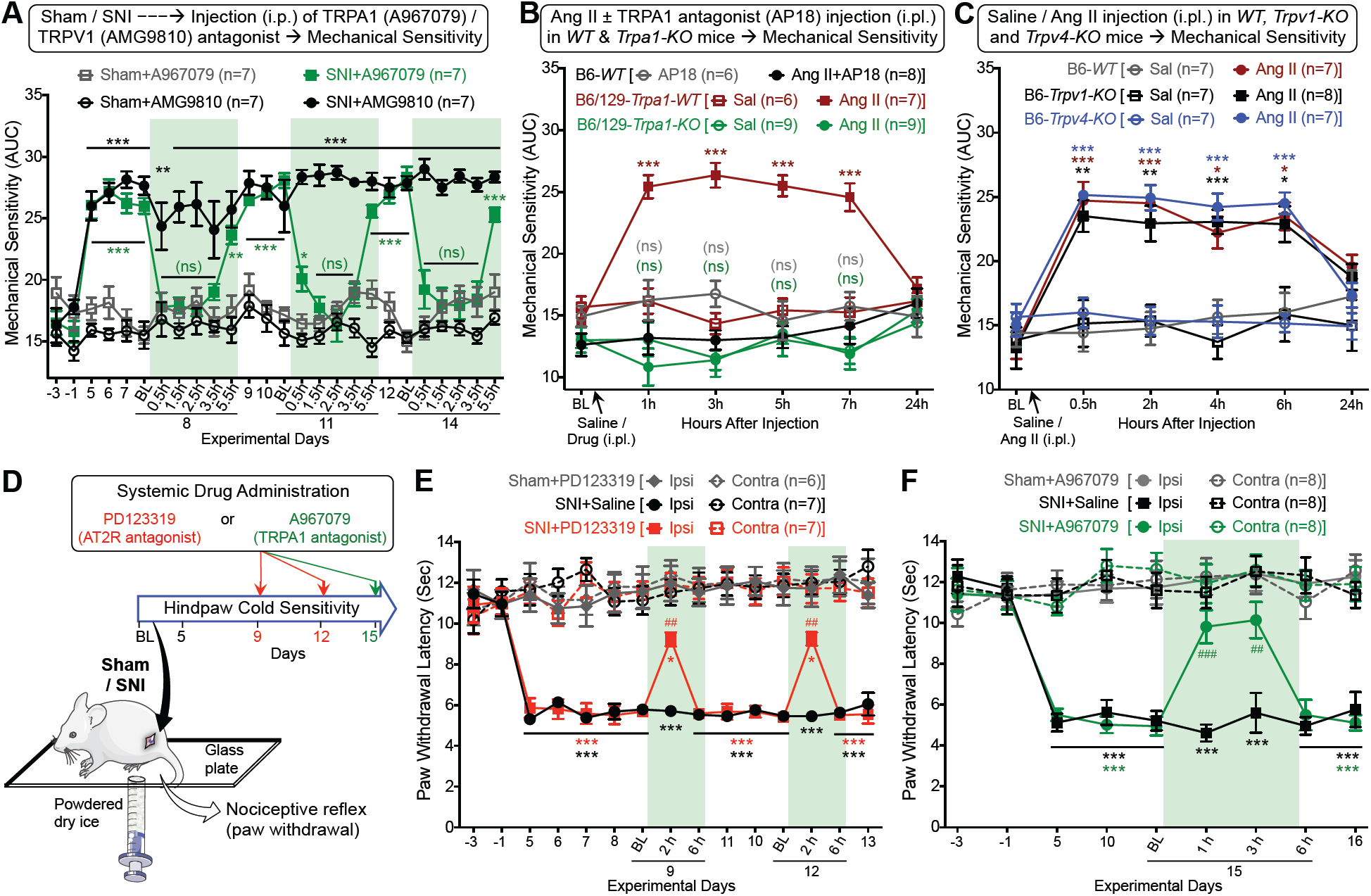
TRPA1 is critical for neuropathic mechanical and cold hypersensitivity. (**A**) TRPA1 antagonist A967079, but not TRPV1 antagonist AMG9810 (30 mg/kg for each; i.p.) attenuates SNI-induced mechanical hypersensitivity in mice. No significant change in mechanical sensitivity is observed in contralateral hindpaws. Mean ± SEM; ^*^*p*<0.05, ^**^*p*<0.01 and ^***^*p*<0.001, versus sham-A967079/AMG9810 groups; not significant (ns), versus sham-A967079 group. (**B**) Co-administration of TRPA1 antagonist AP18 (10 nmol; i.pl.) prevents Ang II-induced (100 pmol, i.pl.) mechanical hypersensitivity in B6-*WT* mice. Ang II (100 pmol, i.pl.) elicits mechanical hypersensitivity in B6/129S-*Trpa1-WT* mice, to a similar extent as seen in B6-*WT* mice, which is absent in B6/129S-*Trpa1-KO* mice. Mean ± SEM; ^***^*p*<0.001, versus B6/129S-*Trpa1-WT*-saline group; not significant (ns), versus B6-*WT*-AP18 and B6/129S-*Trpa1-KO*-saline groups. (**C**) Ang II (100 pmol; i.pl.) induces a similar magnitude of mechanical hypersensitivity in the ipsilateral hindpaws of B6-*WT*, B6-*Trpv1-KO* and B6-*Trpv4-KO* mice. Mean ± SEM; ^*^*p*<0.05, ^**^*p*<0.01 and ^***^*p*<0.001, versus saline injection in respective mouse genotype groups. (**D**) Experimental scheme depicting reflexive cold hypersensitivity assessment, and drug administration timeline in mice subjected to sham/SNI surgery. (**E-F**) Systemic administration of PD123319 (10 mg/kg; i.p.; **E**) and A967079 (30 mg/kg; i.p.; **F**) attenuate SNI-induced cold hypersensitivity. Mean ± SEM; \**p*<0.05 and ^***^*p*<0.001, versus sham+PD123319 or sham+A967079 groups; ^##^*p*<0.01 and ^###^*p*<0.001, versus SNI+saline group. Rectangular boxes denote post-drug administration behavioral assessment time points.

**Figure 4:**
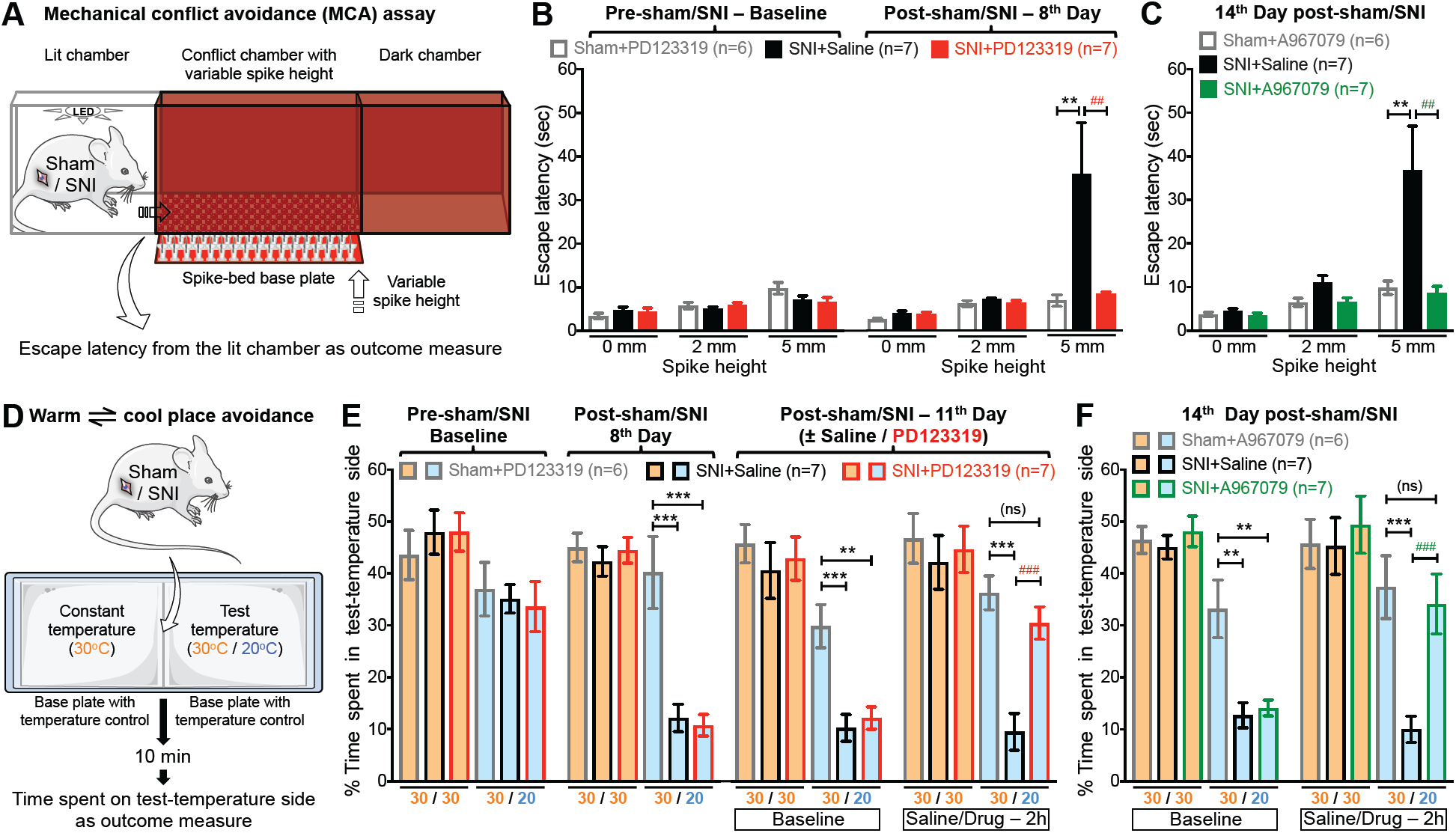
AT2R and TRPA1 are critical for SNI-induced voluntary pain behaviors in mice. (**A**) Experimental scheme depicting ongoing/non-reflexive mechanical hypersensitivity assessment using mechanical conflict avoidance (MCA) system in mice that are subjected to sham/SNI surgery. (**B-C**) Systemic administration of PD123319 (10 mg/kg; i.p.; **B**) and A967079 (30 mg/kg; i.p.; **C**) attenuate SNI-induced increased escape latency from lit chamber to mechanical conflict chamber. Mean ± SEM; ^**^*p*<0.01, versus sham+PD123319 or sham+A967079 groups; ^##^*p*<0.01, versus SNI+saline group. (**D**) Experimental scheme depicting ongoing/non-reflexive cold hypersensitivity assessment using warm/cool plate place avoidance system in mice that are subjected to sham/SNI surgery. (**E-F**) Systemic administration of PD123319 (10 mg/kg; i.p.; **E**) and A967079 (30 mg/kg; i.p.; **F**) attenuate SNI-induced avoidance to cool-temperature chamber. Mean ± SEM; not significant (ns), ^**^*p*<0.01 and ^***^*p*<0.001, versus sham+PD123319 or sham+A967079 groups; ^###^*p*<0.001, versus SNI+saline group.

### Ang II has no direct influence on sensory neuron function

We next investigated how Ang II – AT2R activation in sensory neurons might influence TRPA1 channel activation and/or modulation, which could lead to mechanical and cold hypersensitivity. Prolonged exposure of Ang II (100 nM; 1h) did not elevate intracellular Ca^2+^ ([Ca^2+^]_i_) in cultured mouse or human dorsal root ganglia (DRG) sensory neurons, irrespective of functional TRPA1 expression (Figure 5A, and Supplemental Figure 6 A and B), suggesting Ang II does not directly activate TRPA1 or other TRP channels in DRG neurons. Previous studies have reported that AT2R activation in mouse DRG neurons modulates TRPV1 function (4, 10). Intriguingly, in our experiments Ang II exposure did not influence the function of TRPA1 or TRPV1 channels in mouse or human DRG neurons (Figure 5 B and C). Furthermore, Ang II exposure did not alter action potential (AP) firing, and other properties of membrane excitability of cultured mouse and human DRG neurons (Figure 5D, and Supplemental Figure 6 C and D). Altogether, these findings suggest Ang II does not directly influence sensory neuron function or excitability.

**Figure 5:**
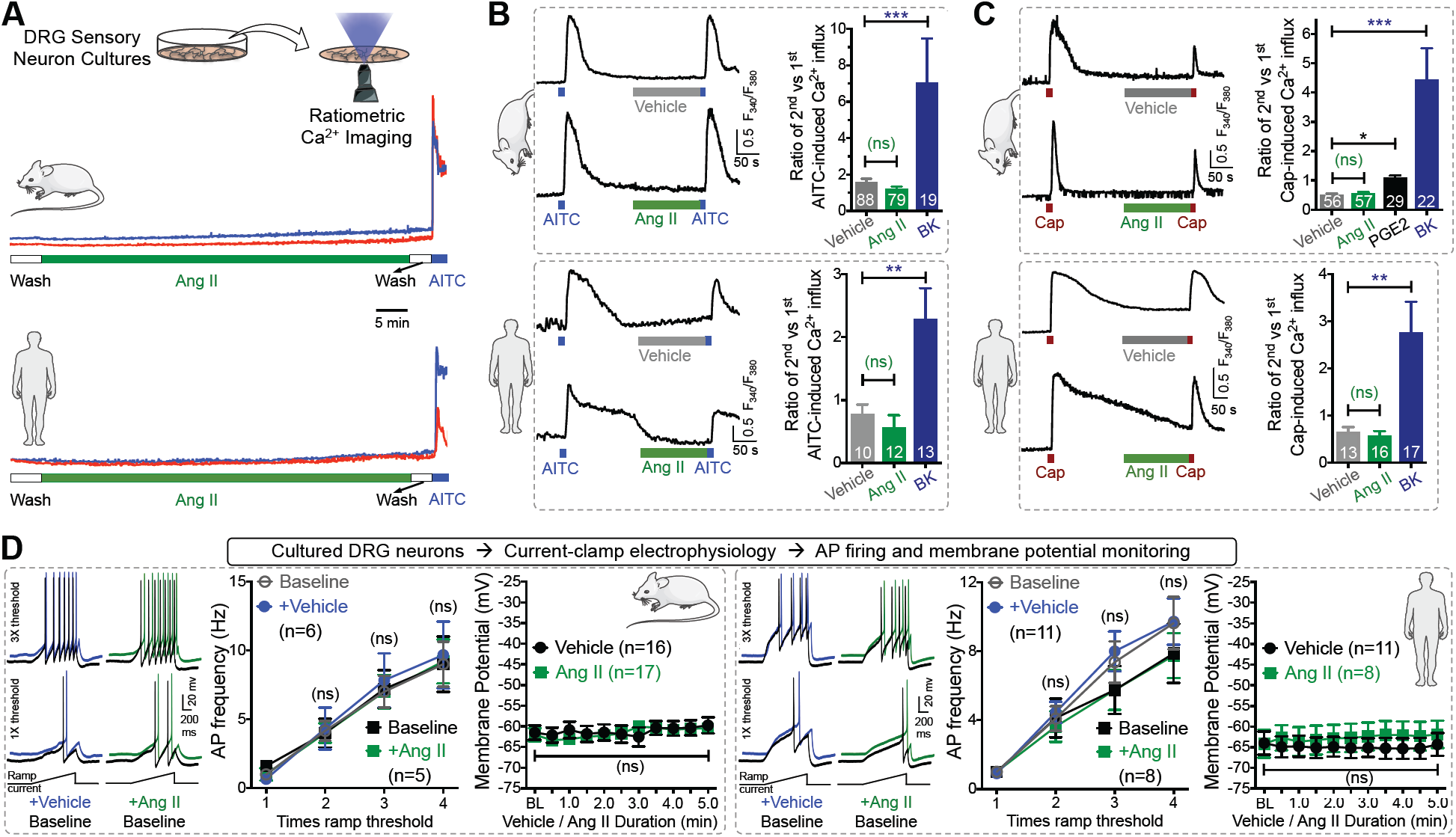
No direct action of Ang II on sensory neuron TRP channels and excitability. (**A**) Ang II (1 μM) induces no [Ca^2+^]_i_ elevation in mouse and human DRG sensory neurons in culture. AITC (100 μM) is used for the detection of TRPA1^+^ neurons. (**B-C**) Ang II (1 μM) fails to potentiate AITC-(50 μM; 15 sec; **B**) and capsaicin (50 nM; 15 sec; **C**) induced [Ca^2+^]_i_ elevation in cultured mouse/human DRG neurons. As positive controls, bradykinin (100 nM) significantly potentiates AITC- and capsaicin-evoked Ca^2+^ flux, and PGE2 (10 μM) potentiates capsaicin-induced Ca^2+^ elevation. Mean ± SEM; \**p*<0.05, ^**^*p*<0.01, ^***^*p*<0.001 and not significant (ns), versus respective vehicle groups. (**D**) Ang II (1 μM; 5 min) fails to influence action potential (AP) firing and membrane potential of mouse and human DRG neurons. Mean ± SEM. AP firing traces for vehicle- and Ang II-treated conditions are offset for distinct visualization from their respective control traces.

### Lack of AT2R expression in DRG sensory neurons

Lack of any direct influence of Ang II on DRG neurons next prompted us to seek evidence in support of functional AT2R expression therein. Ang II exposure did not elicit AT2R-dependent ERK1/2 and p38 MAPK phosphorylation in either mouse or human DRG neurons (Figure 6A, and Supplemental Figure 7), in contrast to previous reports (10, 20). Furthermore, routinely used anti-AT2R antibodies stained DRG tissue from *Agtr2-WT* and *Agtr2-KO* mice with similar intensity (Supplemental Figure 8A), indicating non-specific binding of these antibodies, as has been demonstrated earlier (21). With no credible evidence for AT2R protein expression in DRG neurons, we next made an in-depth investigation of AT2R gene (*Agtr2*) expression in DRG neurons. Immunostaining of DRG sections from C57BL/6-*Agtr2^GFP^* reporter mice, where GFP expression is driven by the *Agtr2* promoter (8), displayed no detectable GFP signal, similar to C57BL/6-*WT* negative controls (Figure 6B, and Supplemental Figure 8B). In the sciatic nerves of *Agtr2^GFP^* reporter mice GFP staining is observed in a subset of NF200^+^ (myelinated) fibers, but not in CGRP^+^ (peptidergic nociceptive neuron marker) fibers (Supplemental Figure 8C). In accordance with this observation, no GFP signal is observed in the nerve fibers in superficial laminae of the spinal cord from *Agtr2^GFP^* mice, where CGRP is expressed by central terminals of sensory neurons (Supplemental Figure 8D). Numerous NF200 and NeuN-positive somata in deeper laminae and ventral horn express GFP (Supplemental Figure 8D), indicating that a subset of central neurons do express AT2R. In particular, GFP expression on ventral horn neurons with larger somata, an anatomical feature of motor neurons, coincide with the expression of GFP in a subset of NF200^+^ fibers in the sciatic nerves (22, 23). Furthermore, no amplification *of Agtr2* mRNA from mouse and human DRGs could be obtained using species-specific AT2R primer sets in qualitative RT-PCR (Supplemental Figure 8E). We next verified high-throughput RNA sequencing data available from published studies and found that *Agtr2* mRNA was either absent or its levels were negligible (below baseline noise values) in mouse DRGs (Supplemental Figure 8F). We also performed deep sequencing on total RNA isolated from human DRGs, and found no significant expression of *AGTR2* mRNA, in contrast to other pain-related channels, receptors and neuropeptide genes (Figure 6C). Complementary to our observations, RNAseq datasets for human DRGs obtained by an independent research group also showed no significant expression of *AGTR2* mRNA in donors without a history of pain or with a medical history of chronic pain conditions (Figure 6D). Interestingly, in both human DRG RNAseq datasets, as well as in previously published mouse DRG RNAseq datasets, significant expression of angiotensinogen mRNA was detected (Figure 6 C and D, and Supplemental Figure 8F). However, renin mRNA expression was undetectable in human and mouse DRGs. Collectively, our in-depth analysis argues against the existence of direct AT2R-TRPA1 signaling in sensory neurons, and suggests that non-neuronal AT2R is involved in Ang II-induced pain hypersensitivity.

**Figure 6:**
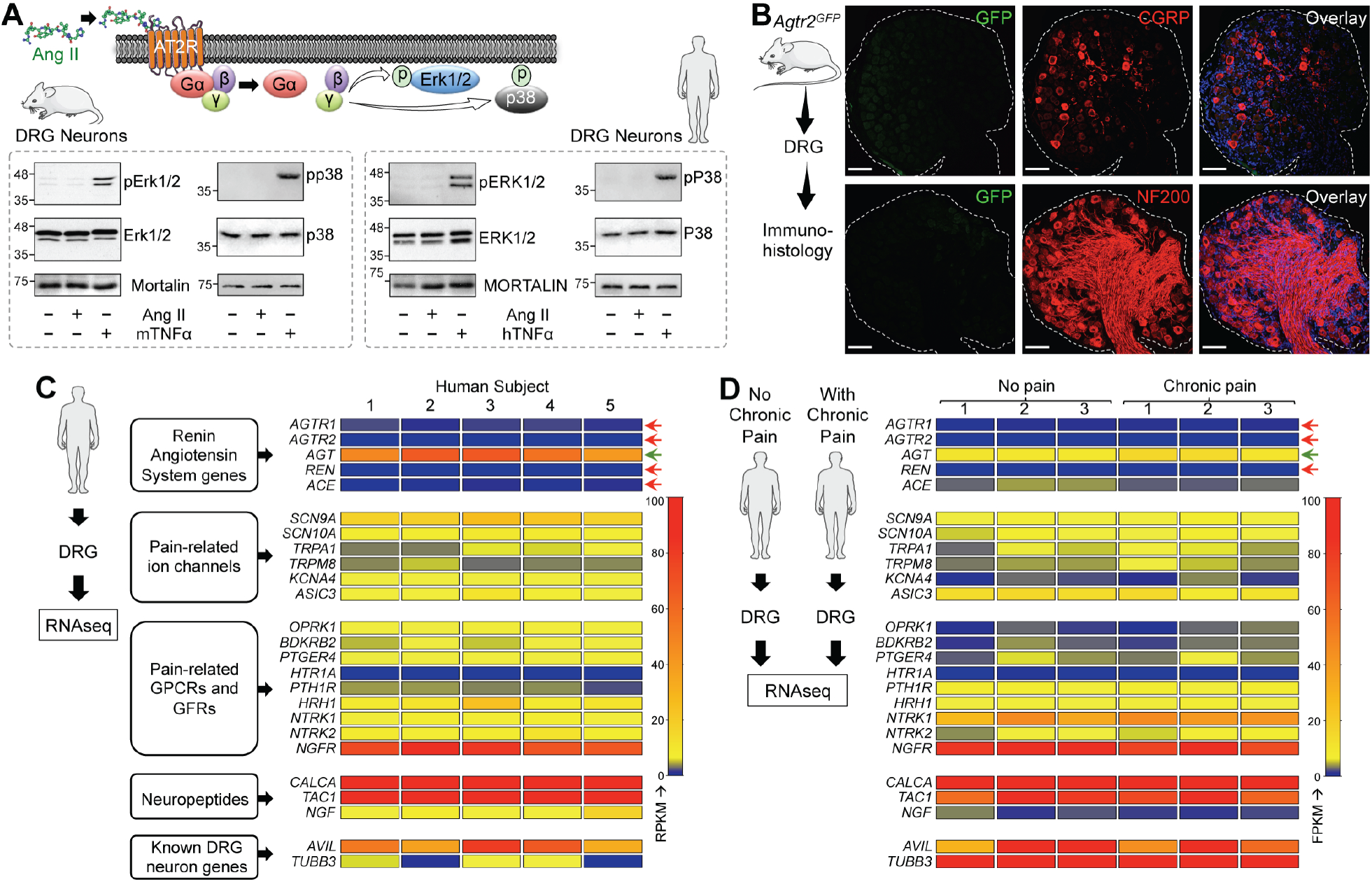
AT2R is not expressed in DRG sensory neurons. (**A**) Ang II (1 μM; 30 min) does not induce phosphorylation of ERK1/2 and p38 MAPK, indicative of Ang II-AT2R activation, in mouse and human DRG neurons. TNF-α (10 nM) is used as a positive control for induction of ERK1/2 and p38 MAPK phosphorylation. (**B**) The *Agtr2* gene (for AT2R) is not expressed in neurons and non-neuronal cells in mouse DRG, as verified by lack of GFP signal in DRG sections from *Agtr2^GFP^* reporter mice, in which the *Agtr2* promoter drives the expression of GFP. DRG sections are stained with CGRP and NF200 antibodies to mark peptidergic and myelinated sensory neurons. Scale bar: 50 μm. (**C-D**) Heat map showing mRNA expression levels (from RNAseq experiments) of renin-angiotensin system (RAS) genes, in comparison to critical pain-associated genes in human DRG tissue (**G**). No alteration in the mRNA levels of RAS genes can be observed in DRGs obtained from humans without or with chronic pain conditions. Red arrows indicate no reliable mRNA expression levels, and green arrow indicates considerable mRNA expression of RAS genes.

### Peripheral macrophages and AT2R expression therein are critical for neuropathic pain hypersensitivity

With no indication of sensory neuron expression of AT2R, or signaling crosstalk with TRPA1 channels within sensory neurons, we investigated the sites of nerve injury and Ang II injection to obtain histological evidence for the underlying mechanism (Figure 7A). SNI induced massive and sustained infiltration of macrophages (MΦs) in both male and female mice, and increased neutrophil infiltration into the site of nerve injury (Figure 7 B to D, and Supplemental Figure 9). Interestingly, the spared sural nerve fibers, which did not show any loss of nerve fiber staining (with NF200), did not show any visible MΦ infiltration. As has been shown previously (24), increased microglial density was observed in the ipsilateral spinal cord dorsal horn of SNI mice, without any detectable neutrophil staining (Supplemental Figure 10). Increased MΦ and neutrophil infiltration was observed in the plantar region of Ang II-injected mouse hindpaws (Figure 7E). Similarly, significant increases in MΦ density are observed in skin biopsies from patients with diabetic neuropathy (DN) and chemotherapy-induced peripheral neuropathy (CIPN) versus healthy controls, a difference associated with loss of PGP9.5^+^ nerve fibers (Figure 7 F and G).

**Figure 7:**
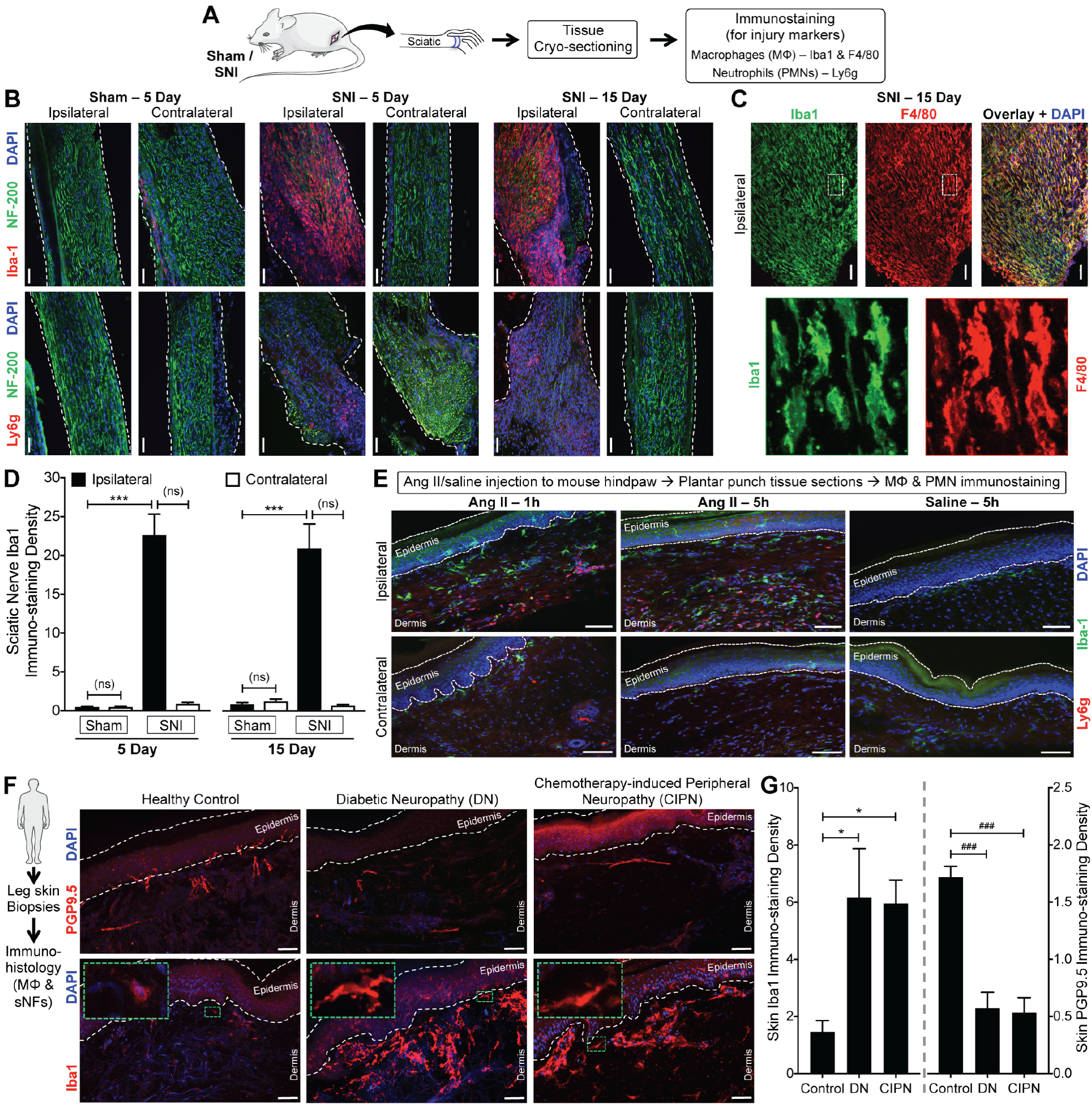
Peripheral macrophage (MΦ) infiltration in mouse and human nerve injury/neuropathy. (**A**) Experimental protocol for identification of injury markers in the sciatic nerve of mice subjected to sham or SNI surgery. (**B-D**) Massive MΦ (Iba1-red, top row in **B**; Iba1-green and F4/80-red in **C**) infiltration and considerable neutrophil (Ly6g-red, bottom row in **B**) infiltration accompany SNI-induced nerve fiber degeneration (decreased NF200 staining; green) in ipsilateral sciatic nerves, 5 and 15 days post-SNI. Sections are co-stained with nuclear marker (DAPI; blue); scale bar: 200 μm. Bottom row images in **C** are magnified view of the area marked with white dotted boxes on top row images. Macrophage density in sciatic nerves is quantified in **D**. Mean ± SEM; ^***^*p*<0.001, versus respective sham-ipsilateral groups; not significant (ns), versus contralateral groups (n= 2 sections per mouse, 4 mice/group). (**E**) Hindpaw Ang II injection (100 pmol i.pl.) enhances MΦ (green – Iba1) and neutrophil (red – Ly6g; blue – DAPI) infiltration both 1 and 5 h post-injection. Scale bar: 100 μm. (**F-G**) Massive MΦ (Iba1-red, DAPI-blue; **F**) infiltration is observed in human leg/ankle skin biopsies from diabetic neuropathy and chemotherapy-induced peripheral neuropathy patients, compared to age-matched healthy controls. This is accompanied by a decrease in the density of nociceptive nerve fibers in the skin (PGP9.5-red, DAPI-blue). Green dotted boxes on the left-top corners in bottom row images represent magnified views of individual macrophages in indicated areas (**F**). Density of both MΦs and nociceptive fibers in skin biopsy are quantified in **G**. Mean ± SEM; ^***^*p*<0.001, versus respective sham-ipsilateral groups; not significant (ns), versus contralateral groups (n= 2 sections each from 8 human subjects/group).

Since AT2R is critical for SNI- and Ang II-induced mechanical and cold hypersensitivity, we next determined if MΦs and/or neutrophils express AT2R. No amplification of AT1R and AT2R genes could be obtained from mouse peritoneal polymorphonuclear neutrophils (PMNs), whereas mRNA for both genes could be amplified from the total mouse peritoneal MΦ RNA (Supplemental Figure 11A). This is also in line with high throughput mRNA expression data on *Agtr2* and other RAS gene expression in mouse and human MΦs available in NCBI-GEO database (Supplemental Figure 11B). Similar to our observations in DRGs, the anti-AT2R antibodies stained peritoneal MΦs with similar intensity from both *Agtr2-WT* and *Agtr2-KO* mice (Supplemental Figure 11 C and D), reiterating the non-specific nature of these antibodies. In order to better validate the expression of *Agtr2* gene on MΦs we next performed immunohistological staining of hindpaw plantar punch biopsies from *Agtr2^GFP^* reporter mice, which show co-localization of GFP with the MΦ marker F4/80 (Figure 8A). Consistent with these observations, Ang II stimulated Erk1/2 phosphorylation, a functional read-out of AT2R expression/activation (25, 26), in B6-*WT, Agtr1a-KO* and *Agtr2-WT* mouse peritoneal MΦs, but not in B6-*WT* mouse PMNs or *Agtr2-KO* MΦs (Figure 8 B and C). Furthermore, Ang II-induced Erk1/2 phosphorylation could be mimicked by the AT2R-selective agonist, CGP42112A, and was blocked by the AT2R antagonist PD123319, but not the AT1R antagonist losartan (Figure 8 B and C), altogether suggesting that functional Ang II – AT2R signaling exists in mouse MΦs.

**Figure 8:**
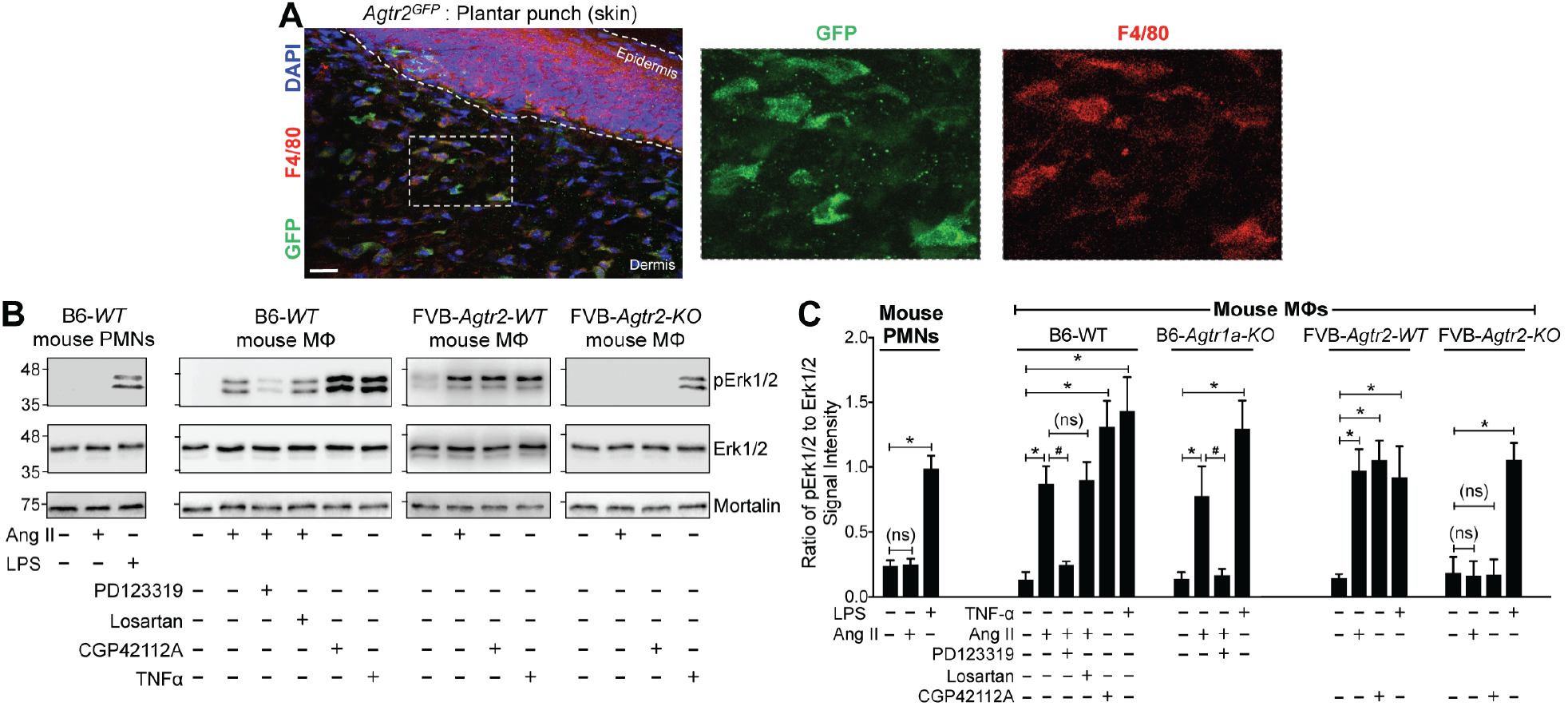
AT2R is expressed in macrophages (MΦs). (**A**) *Agtr2*^GFP^ reporter mouse plantar skin shows co-localization of GFP signal (green) with the MΦ marker F4/80 (red). DAPI: blue; scale: 50μm. (**B**) Ang II (100 nM; 30 min) induces Erk1/2 phosphorylation in mouse (B6-*WT*) peritoneal MΦs, but not in polymorphonuclear neutrophils (PMNs). AT2R inhibitor PD123319 (1 μM), but not AT1R inhibitor losartan (1 μM) attenuates Ang II-induced Erk1/2 phosphorylation in MΦs. Ang II-induced Erk1/2 phosphorylation is absent in MΦs from *FVB-Agtr2-KO* mice, but intact in *FVB-Agtr2-WT* mice. The selective AT2R activator CGP42112A (100 nM) and TNF-α (10 nM) are used in mouse MΦs as positive controls for AT2R activation-signaling and Erk1/2 phosphorylation, respectively. LPS (10 nM) is used as a positive control for Erk1/2 phosphorylation in mouse PMNs. Mortalin (Grp78) immunoreactivity is used as loading control. (**C**) Quantification of the extent of Erk1/2 phosphorylation levels in mouse MΦs and PMNs from experimental groups shown in **B**. Mean ± SEM (n = 3 samples from ≥3 mice per group); \**p*<0.05, ^#^*p*<0.05 and not significant (ns), versus respective comparison groups.

We next verified if MΦs are critical for Ang II-induced pain hypersensitivity. Upon chemogenetic depletion of peripheral MΦs (but not brain and spinal cord MΦs/microglia) with designer drug (B/B-HmD) treatment in *M*acrophage *F*as-*i*nduced *A*poptosis (MaFIA) mice (Figure 9A, and Supplemental Figure 12A), hindpaw injection of Ang II failed to elicit any significant mechanical or thermal hypersensitivity (Figure 9B, and Supplemental Figure 12B). Peripheral MΦ depletion did not lead to gross deficits in peripheral pain perception, as indicated by bradykinin-induced mechanical and heat hypersensitivity in peripheral MΦ-depleted MaFIA mice (Supplemental Figure 12, C and D). SNI in MaFIA mice induced robust mechanical and cold hypersensitivity, similar to that observed in B6-*WT* mice. Furthermore, chemogenetic MΦ depletion in MaFIA mice following SNI progressively attenuated mechanical and cold hypersensitivity, with no influence on heat sensitivity (Figure 9 C to E, and Supplemental Figure 13A). The attenuating effect of MΦ depletion on mechanical hypersensitivity was observed to a similar magnitude in male and female MaFIA mice (Supplemental Figure 13 B and C). As seen earlier in B6-*WT* mice, massive MΦ infiltration was also observed in injured sciatic nerves of MaFIA mice prior to MΦ depletion. In SNI-MaFIA mice, 5 consecutive days of BB-HmD administration (day 11 post-SNI) significantly depleted most peripheral MΦs at the site of injury (Figure 9 C, F and G). Interestingly, recovery from MΦ depletion (day 12 post-SNI onwards) led to progressive re-development of mechanical and cold hypersensitivity, which was accompanied by re-population of infiltrating MΦs in the injured sciatic nerves (Figure 9 F and G). Chemogenetic depletion of MΦs did not influence the increase in MΦs/microglia density in the spinal cord dorsal horn of MaFIA-SNI mice (Supplemental Figure 14).

**Figure 9:**
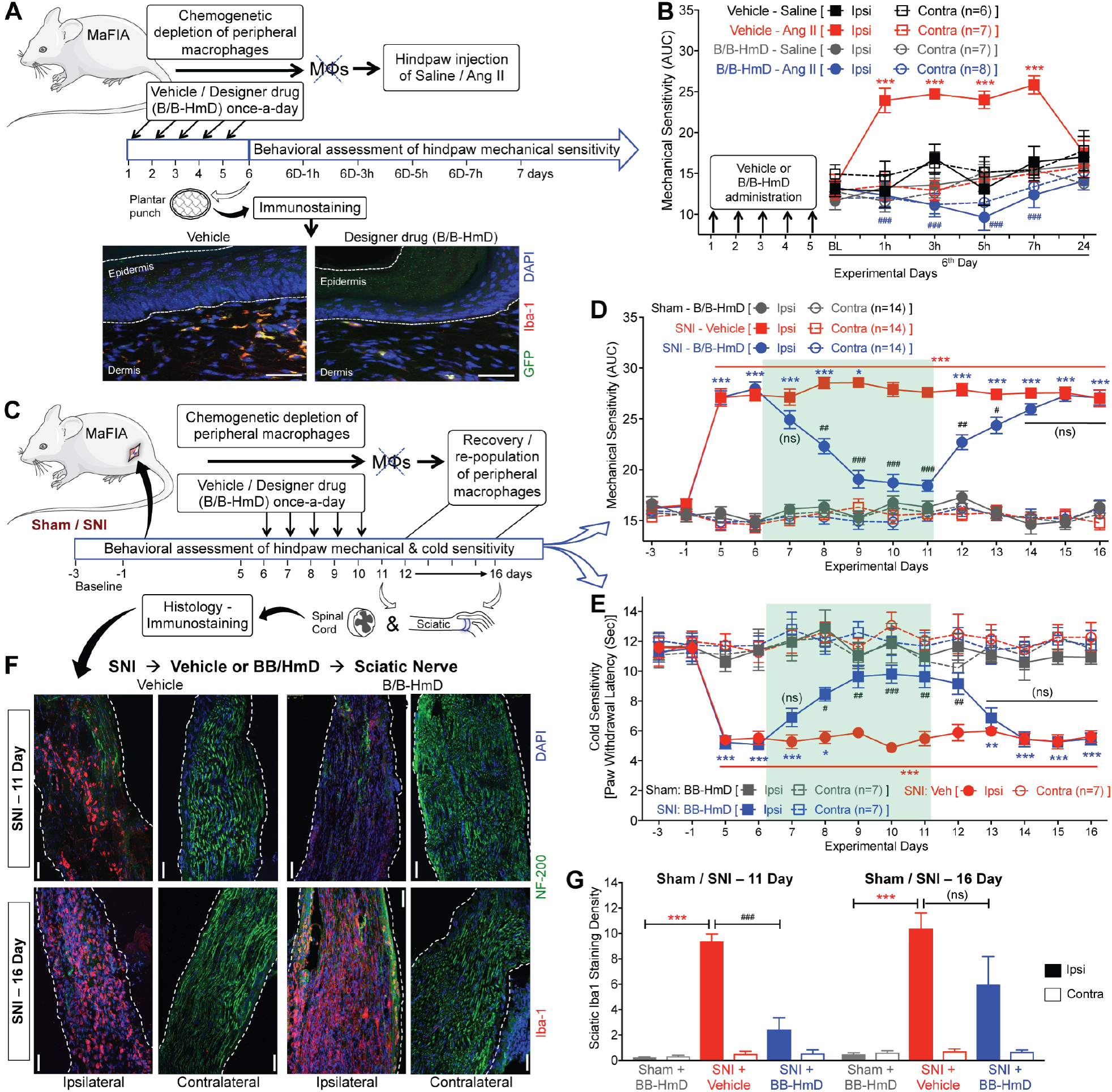
Macrophage (MΦ) infiltration is critical for neuropathic pain hypersensitivity. (**A-B**) Ang II (100 pmol; i.pl.) fails to induce mechanical hypersensitivity in the hindpaws of MaFIA mice subjected to chemogenetic depletion of peripheral MΦs (as shown in a; scale bar: 50 μm) with the administration of a designer drug (B/B-HmD; 2 mg/kg/day for 5 days; **B**). Mean ± SEM; ^***^*p*<0.001, versus vehicle – saline-ipsi group; ^###^*p*<0.001, versus vehicle – Ang II-ipsi group. (**C-E**) Chemogenetic depletion of peripheral MΦs in MaFIA mice with B/B-HmD administration (2 mg/kg/day for 5 days, starting 6 days post-SNI; **C**), leads to significant attenuation of SNI-induced mechanical (**D**) and cold hypersensitivity (**E**), which subsequently returns to pre-depletion levels 3-4 days after the last B/B-HmD administration. Mean ± SEM; \**p*<0.05, ^**^*p*<0.01 and ^***^*p*<0.001 versus sham – B/B-HmD-ipsi group; ^#^*p*<0.05, ^##^*p*<0.01, ^###^*p*<0.001 and not significant (ns), versus SNI – vehicle-ipsi group. (**F-G**) Histological confirmation of MΦ (Iba1-red) depletion at day 11 post-SNI (after 5^th^ B/B-HmD), and re-population at day 16 post-SNI (5 days after final B/B-HmD) in the sciatic nerves (NF200-green) of MaFIA mice (**F**), which are quantified in **G**. Mean ± SEM; ^***^*p*<0.001 versus respective sham – B/B-HmD-ipsi group; ^###^*p*<0.001 and not significant (ns), versus respective SNI – vehicle-ipsi groups (n= 2 sections per mouse, 4 mice/group). Rectangular boxes in panels **D-E** denote post-drug administration behavioral assessment time points.

In order to further test whether the requirement for functional AT2R signaling resides within the immune system, we depleted endogenous hematopoietic progenitor cells by irradiating *Agtr2-WT* recipients and transplanted bone marrow hematopoietic progenitor cells from *Agtr2-WT* or *Agtr2-KO* donors 8 weeks prior to SNI (Figure 10A). *Agtr2-WT* chimeras display similar mechanical and cold hypersensitivity to that seen in B6-*WT* and non-chimeric *Agtr2-WT* mice, which retained responsiveness to treatment with PD123319 on post-operative day 11 (Figure 10 B and C, and Supplemental Figure 15). However, significant attenuation of mechanical and cold hypersensitivity was observed in *Agtr2-KO* chimeras (Figure 10 B and C, and Supplemental Figure 15), despite a similar increase in MΦ infiltration into the sciatic nerve and elevation of microglial density in the spinal cord of *Agtr2-WT* and *Agtr2-KO* chimeras (Supplemental Figure 16). Altogether, these observations suggest that peripheral MΦ infiltration and AT2R signaling therein are necessary for SNI/Ang II-induced pain hypersensitivity.

**Figure 10:**
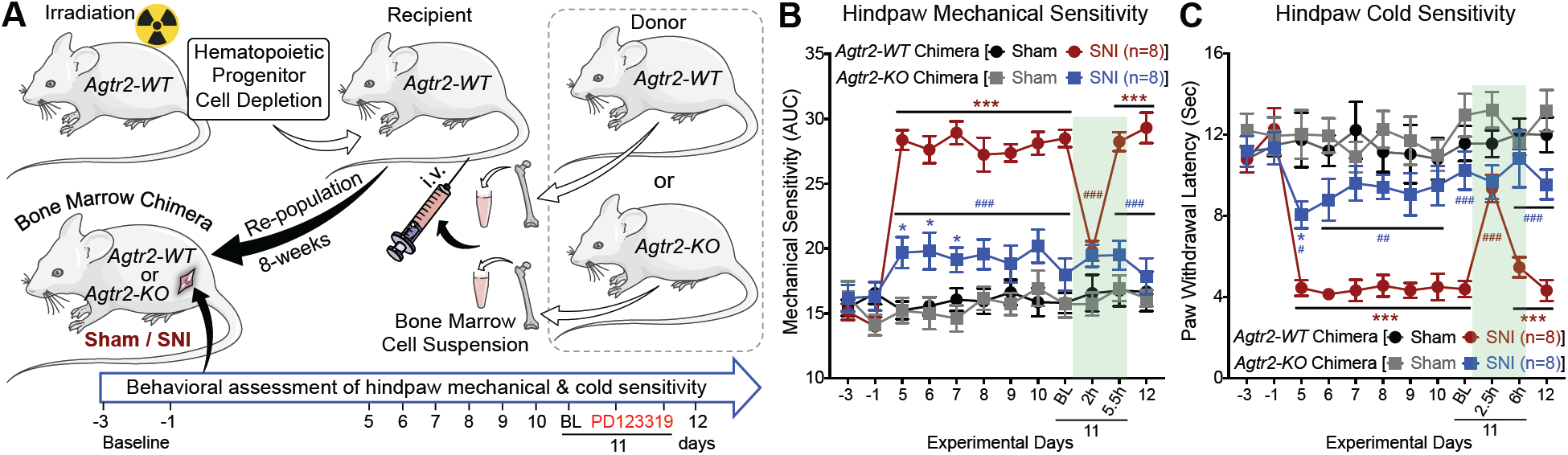
Immune cell AT2R expression is critical for neuropathic pain hypersensitivity. (**A**) Schematic showing generation of *Agtr2-WT* and *Agtr2-KO* chimera mice by bone marrow transplantation, and subsequent induction of nerve-injury/neuropathy for pain-related behavioral assessment. (**B-C**) SNI induces significant mechanical (**B**) and cold hypersensitivity (**C**) in *Agtr2-WT* chimera mice, which could be attenuated by systemic administration of the AT2R antagonist PD123319 (10 mg/kg; i.p.). In contrast, *Agtr2-KO* chimera mice show significantly attenuated mechanical (**I**) and cold hypersensitivity (**J**) upon SNI induction, indicating the critical role of MΦ AT2R in the induction and maintenance of neuropathic pain hypersensitivity. Mean ± SEM; \**p*<0.05 and ^***^*p*<0.001 versus *Agtr2-WT* or *Agtr2-KO* sham-ipsi groups; ^#^*p*<0.05, ^##^*p*<0.01 and ^###^*p*<0.001, versus *Agtr2-WT* SNI-ipsi group. Rectangular boxes denote post-drug administration behavioral assessment time points.

### Macrophage AT2R-mediated redox activation of TRPA1 in sensory neurons

Since MΦs, AT2R and TRPA1 are all required for Ang II-induced peripheral mechanical and cold hypersensitivity, we next investigated the signaling crosstalk between MΦs and sensory neurons. Live cell imaging of mouse peritoneal MΦs showed a time-and dose-dependent increase in DCFDA fluorescence emission [an indication of increased reactive oxygen/nitrogen species (ROS/RNS) production] when exposed to Ang II and Ang III, but not Ang IV (Figure 11 A and B). These observations are in accordance with the reported selectivity and affinity of Ang III for AT2R, as well as the ∼100-fold lower relative affinity of Ang IV for AT2R (27). Contrarily, MΦs did not exhibit any [Ca^2+^]_i_ elevation upon exposure to the TRPA1 activator AITC, before or after Ang II exposure (Supplemental Figure 17), suggesting no functional TRPA1 expression in MΦs. Furthermore, in primary cultures of mouse and human DRG neurons, Ang II exposure did not elicit ROS/RNS production (Supplemental Figure 18A), which is consistent with our failure to detect AT2R expression in DRG neurons. Ang II-induced ROS/RNS production in mouse MΦs could be replicated by the AT2R-selective agonist, CGP42112A, and blocked by the AT2R antagonist PD123319 and the free radical scavenger n-acetylcysteine (NAC), but not by the AT1R antagonist losartan (Figure 11C). ROS/RNS production in mouse MΦs induced by higher concentrations of Ang II and Ang III could also be blocked by PD123319 (Supplemental Figure 18B). No sex-specific differences were observed for Ang II-induced ROS/RNS production in MΦs (Supplemental Figure 18C). Ang II elicited ROS/RNS production in MΦs from *Agtr1a-KO* mice, which could be blocked by PD123319 (Figure 11D). Ang II and CGP42112A exposure also led to elevated ROS/RNS production in MΦs from *Agtr2-WT*, but not from *Agtr2-KO* mice (Figure 11D). Furthermore, we confirmed that similar Ang II/AT2R-dependent ROS/RNS production occurred in MΦs derived from the *Agtr2^GFP^* reporter mouse and the *Agtr2-WT* chimera controls, but not the *Agtr2-KO* chimeras (Figure 11E).

**Figure 11:**
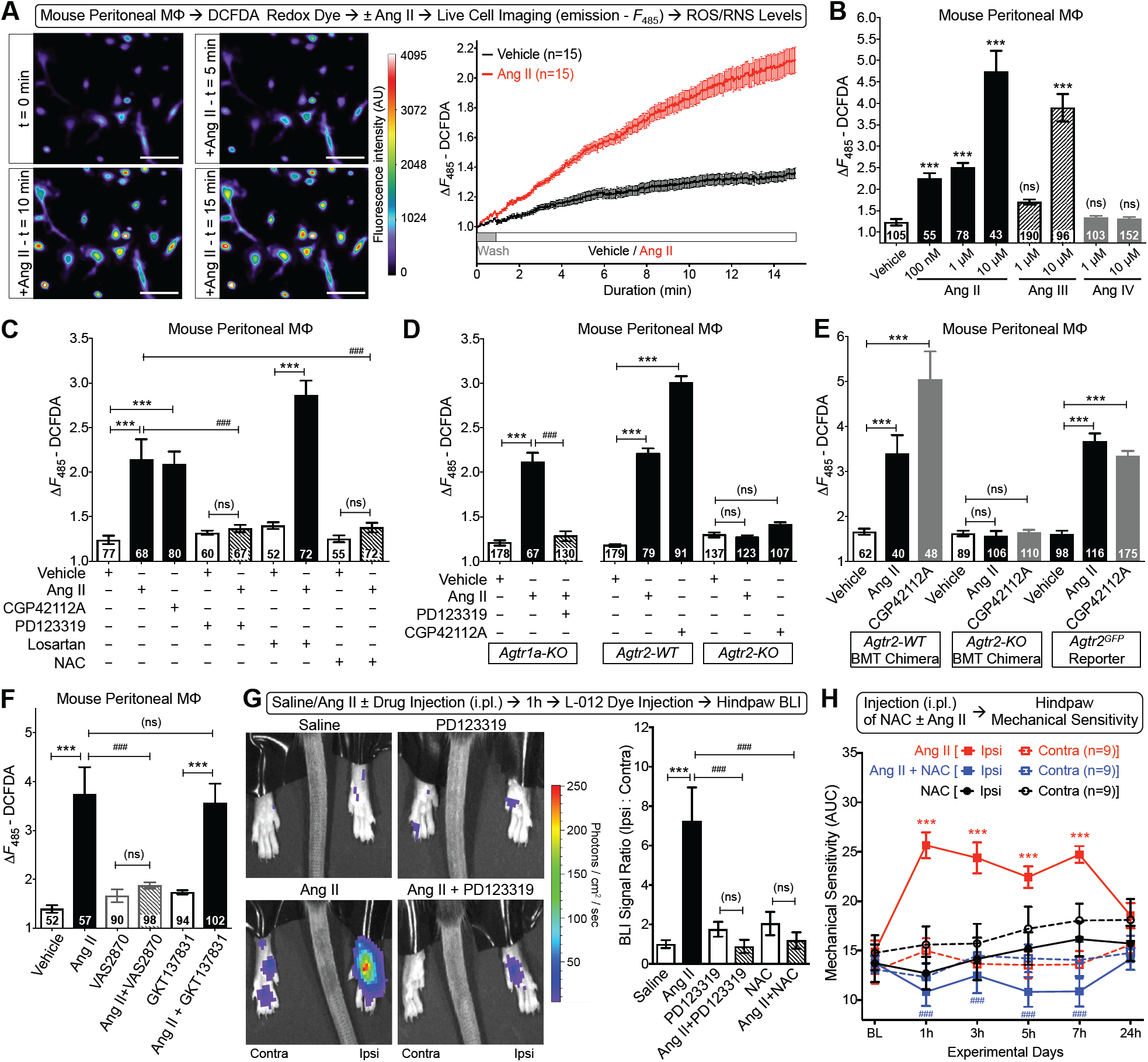
Macrophage (MΦ) Ang II-AT2R redox signaling induces mechanical hypersensitivity. (**A**) Time-lapse images (left) and quantification traces (right) of cultured mouse peritoneal MΦs, showing Ang II-induced (100 nM) elevated ROS/RNS production, as determined by increased intensity of DCFDA redox-sensitive fluorescent dye. Scale bar: 50 μm. (**B**) Ang II and Ang III, but not Ang IV exposure (15 min) dose-dependently induce ROS/RNS production in mouse MΦs. ^***^*p*<0.001 and not significant (ns), versus vehicle group. (**C**) Ang II-induced (100 nM; 15 min) MΦ ROS/RNS production can be attenuated by PD123319 (1 μM) and n-acetylcysteine (NAC; 3 mM), but not losartan (1 μM) co-application. AT2R-selective agonist CGP42112A (100 nM; 15 min) also elevates ROS/RNS levels. Mean ± SEM; ^***^*p*<0.001, ^###^*p*<0.001 and not significant (ns), versus respective comparison groups. (**D**) Ang II (100 nM; 15 min) increases ROS/RNS levels in *Agtr1a-KO* mouse MΦs, which can be attenuated by PD123319 (1 μM) co-application. Both Ang II and CGP42112A (100 nM each; 15 min) increase ROS/RNS levels in FVB-*Agtr2-WT*, but not in FVB-*Agtr2-KO* mouse MΦs. Mean ± SEM; ^***^*p*<0.001, ^###^*p*<0.001 and not significant (ns), versus respective comparison groups. (**E**) Ang II and CGP42112A (100 nM each; 15 min) significantly increase ROS/RNS levels in MΦs from *Agtr2-WT*, but not *Agtr2-KO* chimera mice. Ang II and CGP42112A (100 nM each; 15 min) significantly increase ROS/RNS levels in MΦs from B6-*Agtr2^GFP^* reporter mice, similar to that observed in B6-*WT* mice. Mean ± SEM; ^***^*p*<0.001 and not significant (ns), versus respective comparison groups. (**F**) Ang II-induced (100 nM; 15 min) ROS/RNS production in mouse MΦs can be attenuated by NOX2 inhibitor VAS2870, but not by NOX1/4 inhibitor GKT137831. Mean ± SEM; ^***^*p*<0.001, ^###^*p*<0.001 and not significant (ns), versus respective comparison groups. Numbers in each bar in panels **B-F** indicate date from recorded number of cells in multiple culture batches from ≥4 mice/group. (**G**) Hindpaw injection of Ang II (100 pmol; 1h) increases local ROS/RNS production, as determined by increased L-012 redox-sensitive dye luminescence intensity, and quantified on the graph (right). Co-injection of PD123319 (10 pmol) and NAC (30 nmol) completely attenuate Ang II-induced ROS/RNS production. Mean ± SEM (n=5 mice/group); ^***^*p*<0.001, ^###^*p*<0.001 and not significant (ns), versus respective comparison groups. (**H**) Co-administration of NAC (30 nmol; i.pl.) completely attenuates Ang II-induced (100 pmol, i.pl.) hindpaw mechanical hypersensitivity in mice. Mean ± SEM; ^***^*p*<0.001, versus Ang II – contra, and ^###^*p*<0.001, versus Ang II - ipsi groups.

NADPH oxidase type-2 (NOX2) is the predominant NOX isoform expressed in MΦs, with some expression of NOX1/4 (28). We found that VAS2870, a relatively selective NOX2 inhibitor, largely attenuated Ang II-induced ROS/RNS production MΦs (Figure 11F). However, GKT137831, a selective NOX1/4 inhibitor, did not influence Ang II-induced ROS/RNS production in MΦs (Figure 11F), suggesting AT2R-NOX2 signaling axis as the predominant mechanism. Collectively, these observations suggest that Ang II activation of AT2R in mouse MΦs leads to increased ROS/RNS production, which we next verified *in vivo*. Hindpaw injection of Ang II increased the L-012 luminescence emission intensity, an indicator of increased redox state, which could be attenuated by co-injection of PD123319 or NAC (Figure 11G). Furthermore, co-injection of NAC completely attenuated Ang II-induced mechanical hypersensitivity in mouse hindpaws, without any influence on heat sensitivity (Figure 11H, and Supplemental Figure 19).

Since TRPA1 is a major target for ROS/RNS-mediated sensory neuron excitation (29), we next verified if Ang II-induced ROS/RNS production in MΦs could trans-activate TRPA1 on sensory neurons, using co-cultures of mouse DRG neurons and MΦs. Ang II exposure led to a sustained [Ca^2+^]_i_ elevation in mouse DRG neurons that was dependent on ROS/RNS and TRPA1, and required co-culturing with mouse peritoneal MΦs (Figure 12 A to C). We also verified these effects in co-cultures of mouse DRG neurons with the mouse J774A.1 MΦ cell line, which exhibits functional AT2R expression and Ang II – AT2R-induced ROS/RNS production (Supplemental Figure 20). However, Ang II exposure did not lead to a sustained increase in [Ca^2+^]_i_ in *WT* mouse DRG neurons co-cultured with peritoneal MΦs from *Agtr2-KO* mice (Figure 12C). DRG neurons from *Agtr2-KO* mice exhibited significant modulation of TRPA1 and TRPV1 activation by bradykinin (Supplemental Figure 21), as observed in WT mouse DRG neurons (Figure 5 B and C), suggesting no inherent deficits in TRP channel activation and/or modulation in *Agtr2-KO* DRG neurons. In contrast, Ang II exposure led to a sustained increase in [Ca^2+^]_i_ in *Agtr2-KO* DRG neurons co-cultured with *Agtr2-WT* peritoneal MΦs (Figure 12C), suggesting that functional AT2R signaling in MΦs, but not in DRG neurons, is necessary and sufficient to induce TRPA1 activation in sensory neurons. In accordance with these observations, human DRG neurons co-cultured with the U937 human monocyte-MΦ cell line exhibited similar Ang II/MΦ-dependent increases in [Ca^2+^]_i_, which could be inhibited by the AT2R antagonist PD123319 or TRPA1 inhibitor A967079 (Figure 12D). U937 human monocyte-MΦs exhibit functional AT2R expression, and Ang II-induced ROS/RNS production that is sensitive to PD123319 and NAC, but not to losartan (Supplemental Figure 22). This again confirms that functional AT2R signaling in MΦs, but not in DRG neurons, is necessary and sufficient to induce TRPA1 activation in sensory neurons. Taken together, these results suggest that Ang II activates MΦ AT2R to elicit ROS/RNS production, which then activates TRPA1 on sensory neurons, representing a mechanism of nociceptor excitation that operates in neuropathic conditions (Figure 12E). This signaling axis could be targeted by the AT2R antagonist PD123319 that prevents ROS/RNS production, and by A967079 that blocks TRPA1, to achieve an additive or synergistic relief from nerve injury-induced chronic pain in mice and humans.

**Figure 12:**
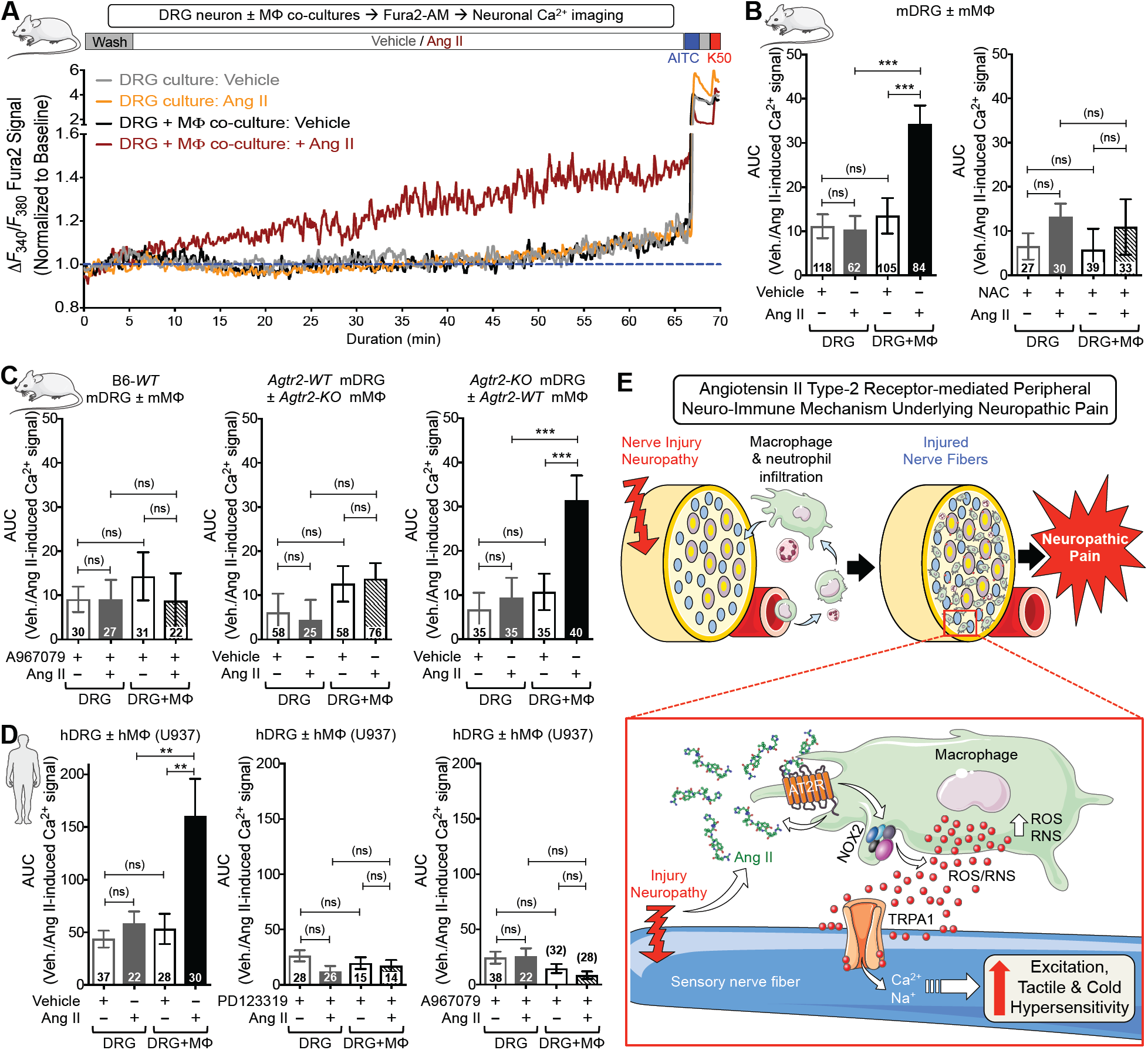
Trans-activation of sensory neuron TRPA1 by Ang II-induced macrophage (MΦ) ROS/RNS production: a mechanism for neuropathic pain. (**A**) Representative traces of Ang II-induced (100 nM, 1h) [Ca^2+^]_i_ elevation in mouse DRG neurons, seen only when co-cultured with mouse MΦs (both B6-*WT* mice). TRPA1^+^ neurons are identified by AITC (100 μM) and 50 mM KCl (K50). Area under the curve (AUC) for Ang II-induced [Ca^2+^]_i_ elevation is subsequently quantified. (**B-C**) Ang II-induced increases in DRG neuron [Ca^2+^]_i_ elevation in co-cultures can be completely attenuated upon co-application of NAC (3 mM; **B**) and TRPA1 antagonist A967079 (1 μM; **C**). Ang II (100 nM, 1 h) fails to induce [Ca^2+^]_i_ elevation in FVB-*Agtr2-WT* DRG neurons co-cultured with FVB-*Agtr2-KO* MΦs; however, increased [Ca^2+^]_i_ remains conserved in FVB-*Agtr2-KO* DRG neurons co-cultured with FVB-*Agtr2-WT* MΦs (**D**). Mean ± SEM; ^***^*p*<0.001 and not significant (ns), versus respective comparison groups (indicated number of cells in culture batches from ≥6 mice/group). (**D**) Ang II (100 nM, 1 h) induces significant elevation in human DRG neuron [Ca^2+^]_i_ levels in co-cultures with U937 human MΦ cell line, which can be completely attenuated upon co-application of AT2R antagonist PD123319 (1 μM) and TRPA1 antagonist A967079 (1 μM). Mean ± SEM; ^**^*p*<0.01 and not significant (ns), versus respective comparison groups (indicated number of cells in culture batches from ≥5 human donors/group). (**E**) The overall model schematic depicts nerve injury/neuropathy leads to infiltration of macrophages and neutrophils, as well as increased local Ang II levels. Ang II activates AT2R receptor on macrophages to induce NOX2-dependent ROS/RNS production, which subsequently activates TRPA1 channel on sensory nerves to enhance neuronal excitation and pain hypersensitivity.

## DISCUSSION

Our findings demonstrate that intercellular crosstalk between macrophages (MΦs) and sensory neurons, mediated by local Ang II-AT2R signaling, constitutes a critical mechanism for chronic pain in nerve injury/neuropathy (Figure 12E). Recent phase II clinical trial findings demonstrate the remarkable analgesic efficacy of the AT2R antagonist EMA401 for PHN-associated neuropathic pain (5). Ang II has been suggested to act directly on DRG neurons to induce neurite outgrowth and PKA-mediated TRPV1 modulation via Gα_s_-coupled AT2R, resulting in peripheral pain sensitization (9, 10). In contrast, activation of Gα_i/o_-coupled AT2R on sensory neurons by a bacterial myolactone toxin has been reported to be analgesic in mice (11). Our findings indicate that AT2R antagonism provides effective analgesia in neuropathic, but not inflammatory pain. Our in-depth investigations show that mouse and human DRG neurons do not express AT2R, nor did we observe any direct influence of sensory neuron function by Ang II. Our study reveals that in nerve injury/neuropathy, elevated local Ang II levels activate AT2R on MΦs to trigger redox-mediated trans-activation of TRPA1 channel on sensory neurons. Uncovering how angiotensin signaling drives neuropathic pain is essential to the understanding of the neurobiological mechanisms and the analgesic effectiveness of AT2R inhibition for neuropathic pain (5).

We demonstrate elevated Ang II levels in injured sciatic nerve, and that the AT2R antagonist dose-dependently attenuates mechanical hypersensitivity induced by nerve injury/neuropathy, but not by chronic hindpaw inflammation. Attenuation of both heat and mechanical hypersensitivity by the same AT2R antagonist in CFA-induced chronic inflammation has been shown previously (30). Our study conclusively shows that Ang II-AT2R signaling and Ang II/nerve injury-induced pain hypersensitivity do not involve TRPV1, the critical transducer of inflammatory heat (and to some extent mechanical) hypersensitivity. Increased infiltration of MΦs and other immune cell types has also been well characterized under CFA-induced inflammatory conditions (31). Although activated immune cells are a source of increased oxidative/nitrosative stress, via multiple mechanisms at the site of inflammation, blockade of AT2R-mediated ROS/RNS production specifically is unlikely to have a measurable analgesic effect in this context. Furthermore, we found no elevation in Ang II levels in CFA-versus saline-injected hindpaws, suggesting a lack of AT2R-induced ROS/RNS production at the site of CFA-induced inflammation, which precludes the effectiveness of an AT2R antagonist for inflammatory pain. With regard to the source of Ang II, MΦs have been shown to express Agt, renin and ACE (32), raising the possibility that the entirety of the RAS required is supplied by MΦs. A scenario where the source of Agt/Ang II is from the liver and/or vasculature, is unlikely, since this would lead to changes in blood pressure, which remains unaltered in SNI animals. Considerable levels of *Agt* mRNA have also been detected in mouse and human DRGs, without any detectable renin mRNA levels, as revealed by RNAseq data. Since renin serves as the first rate-limiting enzyme for the conversion of Agt to Ang II, direct generation and secretion of Ang II by neurons is implausible. It is more likely that sensory nerves secrete Agt, which is then processed by MΦ-derived renin and ACE to produce Ang II upon nerve injury. Further in-depth studies utilizing tissue-specific expression/knock-down of RAS genes are needed to determine the source of Ang II under multiple neuropathies. Furthermore, ROS-mediated oxidation of Agt has been shown to facilitate its enzymatic cleavage by renin to generate Ang I (33). This raises the possibility that Ang II-induced MΦ ROS/RNS production promotes Agt cleavage at the site of nerve injury, in order to establish a feed-forward cycle for the induction and maintenance of neuropathic pain.

Our comprehensive analysis shows no AT2R expression on sensory neurons, clearly implicating non-neuronal AT2R signaling in the development of neuropathic pain. In search of the underlying mechanism, we observed massive MΦ infiltration into the injured nerve, as well as increased density of microglia in the ipsilateral spinal cord dorsal horn, consistent with prior observations (24, 34). We also observed an increase in MΦ density in human skin biopsies from patients with diabetic neuropathy and chemotherapy-induced peripheral neuropathy, which suggests a histological commonality that is conserved in rodent and human neuropathies. Chemogenetic depletion of peripheral MΦs (but not spinal cord microglia) in mice attenuated Ang II- and nerve injury-induced mechanical and cold pain hypersensitivity, indicating MΦs are an indispensable component. Importantly, restoration of mechanical and cold hypersensitivity following re-population of MΦs at the site of nerve injury strongly supports this assertion. Infiltration of MΦs into peripheral nerves and DRG, as well as microglial activation in spinal cord have been implicated in multiple inflammatory, neuropathic and cancer pain conditions. MΦ/microglia-derived inflammatory mediators, growth factors and spinal modulatory signaling have been suggested as the predominant modulatory factors for peripheral pain sensitization (35, 36). Recent findings in a rodent model of experimental trigeminal neuropathy suggest the involvement of MΦ infiltration at the site of nerve injury and increased oxidative stress leads to TRPA1 activation on trigeminal neurons, resulting in pain hypersensitivity (37). This report also shows that systemic chemical-induced depletion of monocyte-MΦs led to attenuation of pain hypersensitivity, with the chemokine CCL2-CCR2 signaling being the critical mediator of monocyte-MΦs and MΦ ROS production (37). However, our chemogenetic approach to specifically deplete peripheral MΦs, and bone marrow transplantation approach to re-populate *Agtr2*-*KO* MΦs in *Agtr2*-*WT* mice precisely identify AT2R function in MΦs as the critical factor for driving this peripheral MΦ-sensory neuron interaction. Furthermore, this interaction is mediated by MΦ AT2R-induced ROS/RNS production, which then trans-activate TRPA1 on sensory neurons to elicit nociceptor excitation. It is important to mention here that potential differences in monocyte-MΦ phenotype and/or differences in specific chemotactic mechanisms operating at sciatic versus infraorbital nerves could result in distinct MΦ-nerve crosstalk mechanisms for DRG versus trigeminal neurons. Our findings on massive MΦ infiltration into injured sciatic nerves in both *Agtr2*-*WT* and *Agtr2*-*KO* chimera mice suggests that AT2R is not required for MΦ infiltration, rather AT2R function on MΦs is critical for driving mechanical and cold hypersensitivity. Therefore, future studies could verify the interdependence of CCL2-CCR2 and AT2R-ROS/RNS signaling in MΦs in relation to specific neuropathic pain conditions.

It has been long understood that central sensitization of peripheral nerve injury responses, which serves as a pain signal amplification system in the CNS, is instrumental in the development of persistent neuropathic pain states (3, 35, 38). Our observations suggest no involvement of spinal cord microglia and/or angiotensin signaling therein; rather, peripheral MΦ AT2R signaling is a critical component that triggers nociceptor excitation and chronic pain in nerve injury/neuropathy. This phenomenon, in combination with spinal cord sensitization pathways, likely drives persistent neuropathic pain states.

Increased expression of RAS components, including AT2R, accompanies the differentiation of MΦs from monocytes (32). Our study comprehensively shows functional expression of AT2R in MΦs, and that its activation by Ang II increases cellular ROS/RNS production. AT2R has been shown to activate NOX (39) that plays a critical role in ROS production, as well as cGMP/nitric oxide synthase (NOS)-mediated production of RNS (40). We found that Ang II-induced ROS/RNS production in MΦs is dependent on NOX2, the predominant MΦ NOX isoform (28). Furthermore, we directly demonstrate increased local AT2R-dependent ROS/RNS production *in vivo* upon Ang II injection. Accordingly, the ROS/RNS scavenger NAC completely attenuates Ang II-induced mechanical hypersensitivity. Prior studies have shown local elevation of H_2_O_2_ levels in injured sciatic nerves in mice, and NOX2-deficient mice exhibit diminished mechanical hypersensitivity in response to peripheral nerve injury (41, 42). Several antioxidants have been proposed as alternative therapeutics for multiple neuropathic pain conditions (43). Importantly, our study demonstrates that local MΦ Ang II-AT2R signaling serves as the critical source of oxidative stress in neuropathic pain. Therefore, AT2R inhibition exerts analgesic effects, presumably via blockade of MΦ-derived ROS/RNS production, which mediates sensory nerve excitation in both mouse and human sensory neurons. Furthermore, oxidative and nitrosative stress have also been shown to induce mitochondrial dysfunction and nerve fiber degeneration (44). Since Ang II-AT2R activation on MΦs is the predominant source of ROS/RNS production, it is plausible that AT2R inhibition could also play a neuroprotective role and thereby contribute in part to its analgesic efficacy.

We observed no sex-specific differences in Ang II-induced ROS/RNS production, or in the induction of mechanical hypersensitivity. Sex differences in the contribution of immune cells, including spinal cord microglia, to chronic pain conditions in mice have recently been detailed (45). That Ang II-AT2R-dependent mechanical and cold hypersensitivity does not appear to involve microglia in the spinal cord may explain the lack of any sex-related differences in peripheral nerve injury-induced pain hypersensitivity.

With regard to how MΦ-derived ROS/RNS act on sensory neurons, our data from SNI and acute Ang II injection experiments pinpoint TRPA1, but not TRPV1 or TRPV4, as the critical nociceptive transducer. Inhibitors of TRPA1, a sensor of mechanical force, cold temperatures and cell damage responses, effectively attenuate mechanical and cold hypersensitivity in rodent models of neuropathy (17, 18, 46, 47). Prior observations have demonstrated that activation of TRPA1 channel at cold temperatures could be potentiated by ROS/oxidative stress, resulting in the development of cold hypersensitivity (17, 18). We show that TRPA1 inhibition, either pharmacologically or with genetic deletion, prevents Ang II/SNI-induced mechanical and cold hypersensitivity. Furthermore, we observed no functional evidence of direct TRPA1 modulation by Ang II in mouse and human DRG sensory neurons, nor did we observe, in contrast to prior observations (11), any Ang II-induced changes in neuronal membrane potential. This may be explained by the different origin of the cells used (PC12 cells and hippocampal neurons (11) versus mouse and human DRG sensory neurons in our studies). Importantly, our study identifies MΦ AT2R-ROS/RNS-mediated activation of neuronal Ca^2+^ influx via TRPA1 as the mechanism underlying intercellular crosstalk between MΦs and DRG neurons. ROS/RNS activate TRPA1 via modification of Cys residues (48, 49), which could serve as the mode of ROS/RNS action on DRG neuron TRPA1. In summary, our study using co-cultures of primary MΦs and DRG neurons shows that AT2R on MΦs and TRPA1 on DRG neurons are critical components of this crosstalk, which is conserved in rodents and humans at a cellular level.

Our findings raise some intriguing possibilities that warrant further exploration. Conditions in which local and/or circulating RAS components are elevated may be associated with mechanical and cold pain hypersensitivity. An association between hypertension and neuropathy has been observed in diabetes mellitus (50, 51). Furthermore, ACE inhibitors have been demonstrated to impact nerve conduction in human diabetic neuropathy (52, 53). Whether this effect could be ascribed to a reduction in Ang II levels and/or Ang II-AT2R-mediated pain sensitization remains unclear, but it may be contingent on MΦ accumulation in the vicinity of damaged nerves. Consistent with this speculation on RAS involvement in pain, our study defines Ang II-AT2R signaling as the indispensable mediator of MΦ-to-sensory neuron crosstalk for the development of chronic neuropathic pain (Figure 12E).

The performance of existing analgesics for neuropathic pain are sub-optimal, and the success rate of new-generation analgesic drug development has been poor (54). This necessitates a comprehensive understanding of the pathophysiology and mechanisms underlying neuropathic pain. With the recent success of an AT2R antagonist, EMA401, for PHN-neuropathic pain (5), our discovery of angiotensin signaling as a critical peripheral sensory nerve-sensitizing mechanism identifies multiple therapeutic targets for neuropathic pain.

## MATERIALS AND METHODS

### Mice

All experiments involving the use of mice and the procedures followed therein were approved by Institutional Animal Studies Committees of Washington University in St. Louis and The University of Iowa, in strict accordance with the *NIH Guide for the Care and Use of Laboratory Animals.* Every effort was made to minimize the number of mice used and their suffering. Mice were maintained on a 12:12 light:dark cycle (06:00 to 18:00 hours) with access to food and water *ad libitum.* Unless otherwise stated, 8 to14 week-old male and female mice were used for all experiments. For details on mouse strains and genotypes used in this study please refer to the Supplemental materials and methods. Specific routes of individual drug injections are mentioned in figures and figure legends. Intraplantar (i.pl.) injections were performed as described previously (55, 56). Mice were manually restrained with the aid of a cloth such that the plantar surface of one hindpaw was exposed. A 10 μL volume was injected into the plantar surface of the hind paw via a 33-gauge stainless steel needle coupled to a Hamilton syringe. Intrathecal (i.t.) injection was performed by lumbar puncture as described previously (57), using a Hamilton syringe and 30-gauge needles to deliver a volume of 5 μL. Peri-sciatic administration of PD123319 or vehicle was performed in a volume of 10μl delivered via a 0.3 ml insulin syringe fitted with a 31-gauge needle. The needle was inserted through the *biceps femoris* muscle, adjacent to the nerve injury incision. Mice were continuously monitored post-injection. Experimenters were blinded to mouse sham/surgery conditions, saline/drug injection types, and injection laterality, as well as to mouse sex and genotypes during the conduct of experiments and data recordings.

### Mouse Spared Nerve Injury (SNI) Model of Neuropathy

Mice underwent surgery as part of the SNI-induced neuropathy, as described previously in several reports (14, 15, 58). See the Supplemental materials and methods for full details.

### Behavioral Assessment of Heat, Cold and Mechanical Hypersensitivity

Heat, cold and mechanical sensitivity on mouse hindpaws were assessed as described in numerous reports (55, 56). See the Supplemental materials and methods for full details.

### Warm-Cool Place Avoidance Assay

Warm-cool place avoidance test was performed as a non-reflexive assessment of ongoing cold hypersensitivity-pain behavior due to neuropathy in mice (18, 59), following published procedures, with minor modifications. See the Supplemental materials and methods for full details.

### Mechanical Conflict Avoidance Assay

Voluntary mechanical conflict avoidance (MCA) in mice was assessed following described protocol, with modifications (19). See the Supplemental materials and methods for full details.

### Primary Cell Culture

Mouse DRG neurons were isolated, dissociated and cultured on coated glass coverslips, as detailed in previous reports (55, 56). Neurons were used within 2-3 days of culturing. Human DRGs from consented donors (3 females, mean age 30 years, 1 month; 6 males, mean age 26 years, 8 months) were acquired through MidAmerica Transplant Services (St. Louis, MO) and prepared as detailed in several recent reports (60, 61). Neurons were used within 5-6 days of culturing *in vitro*. Mouse peritoneal macrophages were isolated as described in the literature (62). Mouse neutrophils were isolated from peritoneal fluid as described previously (63). The human monocyte-MΦ cell line U937 was cultured, and differentiated into MΦs with the treatment of phorbol 12-myristate 13-acetate (PMA) and recombinant human GM-CSF 24 hours prior to use. For details on co-cultures of mouse/human DRG neurons with macrophages please refer to the Supplemental materials and methods.

### Angiotensin II Enzyme immunoassay (EIA)

Quantification of angiotensin II levels in mouse sciatic, spinal cord and plantar tissue were performed by enzyme immunoassay (EIA) using the Ang II EIA kit (Sigma-Aldrich), according to the manufacturer’s instructions. See the Supplemental materials and methods for full details.

### Immunohistochemistry

Brain, spinal cord, DRG, sciatic nerve and plantar punch biopsies were harvested from mice as previously described in numerous reports (64-66). 40 μm-thick fixed frozen sections of mouse brain, spinal cord and plantar punch, 25 μm-thick sections of mouse DRGs and 50 μm-thick sections of human punch biopsies, harvested from the lower leg/ankle region (demographic details listed in Supplemental table 1) were used. See the Supplemental materials and methods for full details, as well as the Supplemental tables 2-3 for details on antibody source and dilutions.

### ImageJ quantification

Density of Iba1 staining in spinal cord and sciatic nerve, and Iba1/PGP9.5 in plantar skin was quantified using ImageJ as described previously (67). Threshold RGB intensity was set in a blinded fashion and maintained between images to be compared. Approximately 1.1 mm of sciatic nerve and 0.5 mm^2^ of skin were captured in each field. The area of the ROI that exhibited fluorescence above threshold was recorded as a percentage value.

### Live Cell Imaging

Functional calcium imaging on DRG neurons and macrophages was performed as described previously (55, 56). Fura 2-AM calcium-sensitive dye was used for calcium imaging, and for the quantification of cellular reactive oxygen/nitrogen species (ROS/RNS) levels 2’,7’-Dichlorofluorescein diacetate (DCFDA) dye was used. See the Supplemental materials and methods for full details.

### RNA Deep Sequencing

Deep sequencing was performed on total RNA isolated from independently obtained human DRG tissue at two sites. Human lumbar DRGs from tissue donors without any history of pain-related disease conditions, and from donors with a history of chronic pain conditions were obtained through MidAmerica Transplant Services (St. Louis, MO) and AnaBios Inc. (San Diego, CA). Demographic details of human DRG tissue donors are provided in Supplemental table 4. See the Supplemental materials and methods for full details.

### RNA Sequencing Data Availability

Reference genomes and transcriptomes used for human RNA-seq mapping were NCBI hg19 and Gencode v14, respectively (PMCID: PMC3431492). Data from RNA-seq experiments for the normal, non-pain donors is submitted to dbGaP with the accession number phs001158.v1.p1. The pain donor samples are in process of submission to dbGaP.

### Western Blotting

Total protein lysates of cultured DRG neurons, MΦs and neutrophils were prepared, as described previously (55, 56). See the Supplemental materials and methods for full details.

### Irradiation and Mouse Bone Marrow Transplantation

FVB-*Agtr2-WT* recipient mice were irradiated and injected with donor marrow harvested from either FVB-*Agtr2-WT* or FVB-*Agtr2-KO* donor mice following standard procedure. See the Supplemental materials and methods for full details.

### Bioluminescence Imaging of Reactive Oxygen/Nitrogen Species **in vivo**

Imaging and quantification of ROS/RNS levels in mouse hindpaws following saline or Ang II injection (i.p.) were performed using the fluorescent ROS/RNS indicator dye L-012 (50 mg/kg) as described previously (68). See the Supplemental materials and methods for full details.

### Electrophysiology

Current-clamp recordings from mouse and human DRG neurons were performed, as described previously (55, 56, 60). See the Supplemental materials and methods for full details.

### Chemicals and Reagents

Details of specific type and source of chemicals and reagents used in this study can be found in the Supplemental materials and methods.

### Statistical Analysis

Data are presented as mean ± SEM. For behavioral experiments, two-way ANOVA with Tukey’s multiple comparisons *post hoc* test was performed. *p* < 0.05 in each set of data comparisons was considered statistically significant. Biochemical, Ca^2+^, DCFDA, and L-012 imaging data were analyzed using one-way ANOVA with Tukey’s or Bonferroni’s multiple comparisons *post hoc* test. Unpaired Student’s *t*-test was used for comparison of experimental results from less than 3 groups. All analysis was performed using GraphPad Prism 7.0 (GraphPad Software, Inc.).

## Acknowledgements

We are thankful to Samantha Kelly and Masato Hoshi for technical assistance, Drs. Justin Grobe, Nicole Littlejohn and Carmen Halabi for help with *Agtr2-WT* and *Agtr2-KO* mouse breeding, and Dr. Mathias Leinders for help with i.v. injections in mice and advice on plasma extravasation assays. Help and assistance provided by Eric Tycksen from the Genome Technology Access Center (GTAC), Washington University in St. Louis (NIH-CA91842 and UL1TR000448), and Andrew Torck and Matthew Neiman (University of Texas, Dallas) in executing the RNA-seq data quantification and analysis are gratefully acknowledged. *Trpv4-KO* mice were generously provided by Dr. Wolfgang Liedtke. *Agtr2-KO* mice (originally generated by Drs. Victor J. Dzau, and Richard E. Pratt) were generously provided by Drs. Curt D. Sigmund and Justin L. Grobe (The University of Iowa Carver College of Medicine). We are grateful to Mid-America Transplant, AnaBios Inc., the families of human DRG donors, and to all the human volunteers (healthy or with neuropathic pain conditions), whose skin biopsies were used in this study. We acknowledge Dr. Joseph Vogel for the generous gift of U937 human macrophage cell line (originally obtained from the ATCC) for this study. The vast majority of this study was supported by funds from the Washington University Pain Center and Washington University School of Medicine, Department of Anesthesiology. Additional funding sources that supported this study are: NIH-grant NS069898 (to D.P.M.); NIH grant CA171927 (to A.D.M.); Danish Diabetes Academy, supported by the Novo Nordisk Foundation (to P.K.); NIH grants DK102520 and U01DK101039 (to S.J.); NIH grants HL125805 (to A.D.dK.); NIH grant NS072432 (to Y.M.U.); NIH grant NS065926 (to T.J.P.); University of Texas STARS funding (to T.J.P. and G.D.); and NIH grant NS42595 (to R.W.G.). We would like to extend our gratitude to Dr. Michael R. Bruchas for suggestions and critical comments on this manuscript; Professors Troels Staehelin and Jens Randel Nyengaard for their input and support on human skin biopsy experiments; and Professors Donna L. Hammond and Alex S. Evers for their continued support and encouragement for this work.

## Author Contributions

A.J.S. and D.P.M. conceived and designed the entire study and majority of the experiments therein. Specific contributions: A.J.S. and D.P.M. performed most mouse behavioral experiments, with contributions from A.D.M. and V.K.S.; A.J.S. and D.P.M. performed the analysis of most mouse behavioral experiments with contributions from A.D.M.; A.J.S. performed all Ca^2+^ and ROS/RNS live-cell imaging experiments and data analysis, with contributions from S.M.T.; B.A.C. performed most electrophysiology experiments on human and mouse sensory neurons, with contributions from A.D.M., and both B.A.C. and A.D.M. performed data analysis; S.K., D.P.M., and A.J.S. performed and analyzed data pertaining to RT-PCR, Western blots and EIA experiments; A.J.S. performed immunohistological experiments and data analysis for mouse tissue, and P.K. performed immunohistological experiments on human skin biopsy tissue, with contributions from A.J.S. and S.H. on data analysis; J.P.G. performed a significant portion of mouse neuropathic surgeries and intrathecal drug administration; M.R.M. and B.S.K. performed mouse bone-marrow transplantations; A.D.dK. and E.G.K. provided the AT2R reporter mice and contributed to immunohistological experiments on tissue from these mice; M.V.V., L.A.M., and T.D.S. contributed to human DRG isolation and cultures, along with contributions from A.J.S.; P.R.R., S.J., G.D., T.J.P., R.W.G. performed and analyzed RNAseq experiments from human DRGs; Y.M.U. contributed to mouse chemogenetic experiments and study design for these experiments; and R.W.G. provided laboratory facilities for electrophysiological and human DRG neuron cultures utilized in this study. A.J.S. and D.P.M. wrote the paper, with input from the rest of the authors.

## List of Supplemental materials

1. Supplemental materials and methods in detail.
2. Supplemental Tables: Supplemental Table 1. Human skin biopsy tissue donor demographic details Supplemental Table 2. Primary antibodies used in this study Supplemental Table 3. Secondary antibodies used in this study Supplemental Table 4. Oligonucleotides used in RT-PCR experiments Supplemental Table 5. Human donor demographic details for RNAseq experiments
3. Supplementary Figures: Supplemental Figure 1: AT2R antagonist dose-dependently attenuates nerve injury-induced mechanical hypersensitivity, without influencing heat hypersensitivity in mice. Supplemental Figure 2: AT2R antagonist does not produce any hemodynamic changes in mice. Supplemental Figure 3: AT1R and AT2R antagonists do not attenuate inflammatory pain hypersensitivity in mice. Supplemental Figure 4: Ang II injection induces AT2R-dependent mechanical, but not heat hypersensitivity in mice. Supplemental Figure 5: Involvement of AT2R and TRPA1 in SNI-induced reflexive and voluntary mechanical and cold hypersensitivity. Supplemental Figure 6: Ang II does not influence [Ca^2+^]_i_ levels and membrane excitability in mouse or human sensory neurons. Supplemental Figure 7: No functional AT2R expression in mouse or human sensory neurons. Supplemental Figure 8: AT2R is expressed in the spinal cord, but not in DRG neurons. Supplemental Figure 9: No sex differences in SNI-induced macrophage infiltration in mouse sciatic nerves. Supplemental Figure 10: SNI increases spinal Iba1 expression in mice. Supplemental Figure 11: Mouse macrophages (MΦs) express AT2R. Supplemental Figure 12: Mouse peripheral macrophages (MΦs) are critical for mechanical, but not heat hypersensitivity. Supplemental Figure 13: Mouse macrophages (MΦs) are critical for SNI-induced mechanical hypersensitivity. Supplemental Figure 14: Chemogenetic depletion of mouse peripheral macrophages (MΦs) does not influence SNI-induced enhanced spinal microglia density. Supplemental Figure 15: *Agtr2* expression within the immune system is required for mechanical hypersensitivity associated with SNI. Supplemental Figure 16: *Agtr2* in immune cells is not required for SNI-induced increased macrophage and microglia density. Supplemental Figure 17: TRPA1-induced calcium influx is absent in macrophages (MΦs). Supplemental Figure 18: Angiotensin receptor signaling is present in macrophages (MΦs), but not DRG neurons. Supplemental Figure 19: N-acetyl cysteine (NAC) does not influence hindpaw heat sensitivity in mice. Supplemental Figure 20: J774A.1 mouse monocyte/macrophage (MΦ) cells express functional AT2R. Supplemental Figure 21: *Agtr2*-*KO* DRG neurons show normal TRP channel activation/modulation. Supplemental Figure 22: U937 human monocyte/macrophage (MΦ) cells express functional AT2R.

